# Spatial deconstruction of the plasma membrane

**DOI:** 10.1101/2025.11.14.688519

**Authors:** Venkata Shiva Mandala, Chen Zhao, Qiuying Chen, Steven Gross, Jacob Geri, Roderick MacKinnon

## Abstract

The plasma membrane of cells is known to be heterogeneous with regards to the spatial distribution of proteins and lipids, but the nature and origins of this heterogeneity are unclear. In this study we perform fluorescence microscopy on plasma membrane sheets and find that most proteins occur in protein-rich domains separated by intervening protein-poor regions. We show that the protein-rich domains and protein-poor regions are at least partially preserved in plasma membrane-derived vesicles that lack cytoskeletal elements, permitting their separation and isolation by density centrifugation. Compositional analysis by nuclear magnetic resonance and mass spectrometry shows that different proteins and lipids exhibit characteristic tendencies to partition into the protein-rich domains. This differential partitioning is correlated with the function of the different proteins, suggesting segregation based on cellular processes. Likewise, ordered lipids including cholesterol and sphingomyelin differentially segregate, being more abundant in the protein-rich domains. We propose that the collective assembly of certain proteins and lipids creates the heterogenous distribution of membrane components and the emergence of protein-rich domains. These domains could create distinct environments for the function and segregation of various membrane processes.

## Introduction

The plasma membrane of a living cell serves not only to insulate the cell from its environment but also as a staging for signaling processes to interact with the outside world ^1^. The organization of this membrane thus underlies many important biological processes, including signal transduction, cell-cell interactions, and nutrient transport. At the simplest level, the plasma membrane is a composite of proteins and lipids that assemble into an organized structure that supports the array of biological functions carried out in the membrane ^2^.

It is well known from microscopy and molecular tracking studies that the distribution of individual proteins and lipids in the plasma membrane is heterogeneous ^3^. Various models have been proposed to explain this heterogeneity – protein-protein interactions ^4,5^, partitioning into membrane domains with specific properties ^6,7^, and segregation by cytoskeletal elements ^8,9^. Yet there is no consensus on the general principles of plasma membrane assembly and their underlying mechanisms ^2^.

We became interested in this topic while studying signaling pathways in cardiomyocytes using immunolabeling. We observed that many membrane proteins – including G-protein coupled receptors, ion channels, and an enzyme – form self-specific clusters, termed higher order transient structures or (HOTS), indicative of weak protein-protein interactions ^10^. These HOTS protein clusters did not appear in all regions of our micrographs but were rather concentrated in regions that appear electron dense in our uranyl-staining experiments. These regions, which we term “protein-rich domains” (and some others have called protein islands ^11–17^) seemingly harbored most of the protein clusters that we studied.

At the same time, using cryo-electron microscopy we were also studying the structures of proteins in vesicles derived directly from the plasma membranes of cells ^18^. We noticed that these vesicles were highly heterogenous in their appearance; some contained a high density of proteins, whereas others appeared relatively empty. We wondered whether the protein-rich vesicles were derived from the protein-rich domains observed in our immunolabeling experiments, and the protein-poor vesicles from the intervening protein-poor regions of the plasma membrane. If this were the case, the cell-derived vesicles could prove useful to isolate and study spatial domains present in the plasma membrane.

Here, the first set of experiments examines the existence of protein-rich domains in cell membranes. We used unroofed cell membranes and mildly fixed whole cells, which are subject to artifacts. We note, however, that others have observed patterns in living cells and in various other plasma membrane preparations that are consistent with those we observe ^10–16^. The second set of experiments establishes the persistence of relatively protein-rich and protein-poor domains in giant plasma membrane vesicles derived from the cell membrane. The third set of experiments analyzes the protein and lipid compositions of fractionated protein-rich and protein-poor vesicles derived from the larger vesicles, and thus from the plasma membrane. Our analysis leads us to conclude that most but not all membrane proteins are concentrated in protein-rich domains in the plasma membrane. A smaller subset of proteins with distinct functions are not concentrated but rather are more evenly distributed over the membrane. Further, lipids are also unevenly distributed, with sphingomyelin and cholesterol existing at higher concentrations in protein-rich domains.

These experiments demonstrate the existence of distinct domains in plasma membranes, how to isolate them, and how to characterize their composition. Our initial analysis shows that functionally connected proteins reside within domains, implying that they subserve membrane function.

## Results

### Protein-rich domains in unroofed membranes and in intact cells

To study the distributions of all membrane-associated proteins in the plasma membrane, we labeled proteins specifically but indiscriminately using a cysteine-reactive fluorophore and labeled lipids using a membrane-partitioning dye (**Fig. 1A**). This targeted, indiscriminate labeling was carried out either on membrane sheets obtained by unroofing the cells or on intact, fixed cells (**Fig. 1**). We used HEK293 cells grown on glass coverslips coated either with poly-D-lysine or both poly-D-lysine and laminin, as these same cells were used later in suspension culture to generate the cell-derived vesicles. Similar domains have been reported previously in cardiomyocytes, T cells, B cells, mast cells, PC12 cells, and in COS-7 cells grown on different substrates ^10–16^.

**Figure 1.**
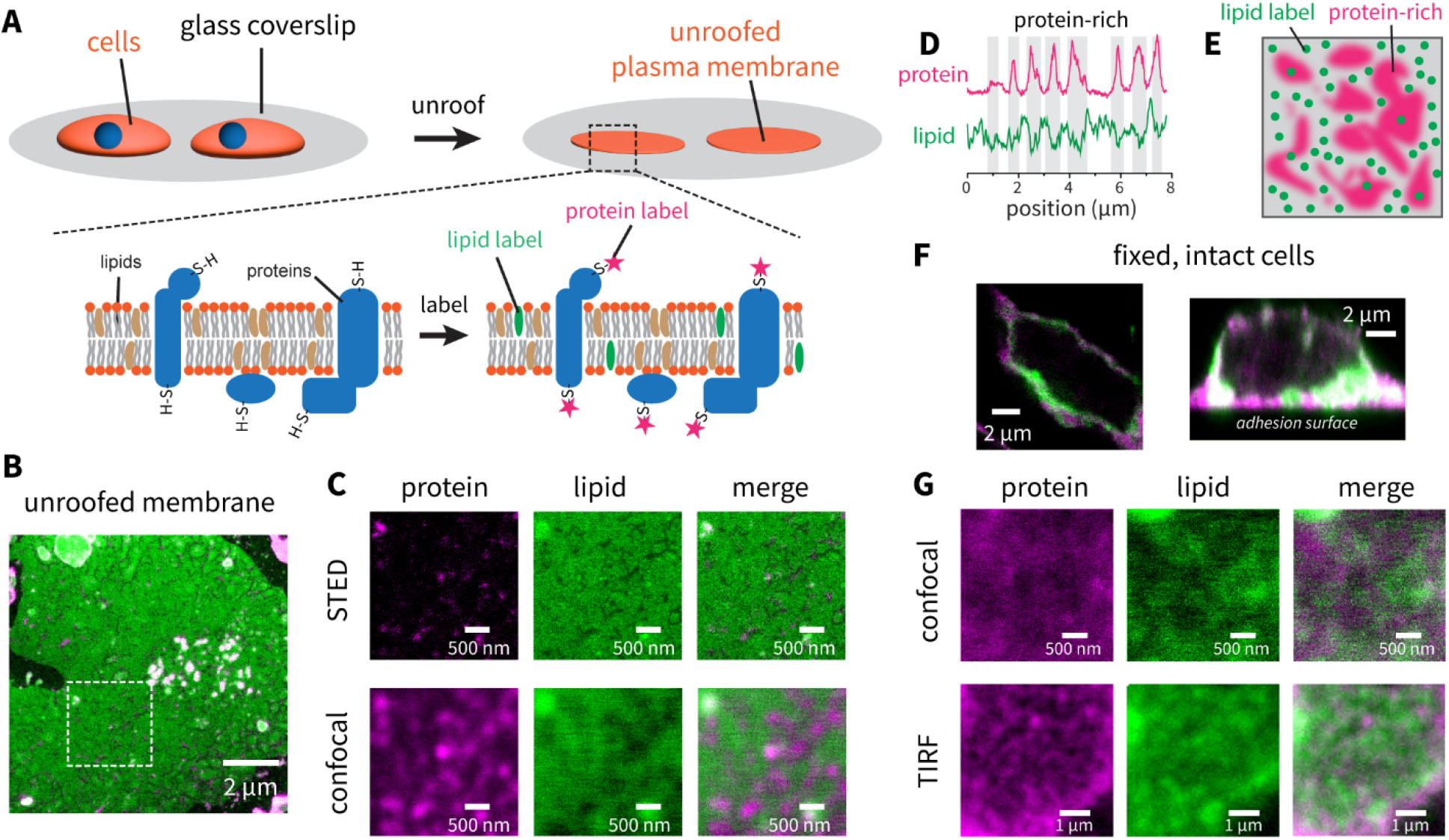
Labeling of proteins and lipids in plasma membranes reveals protein-rich domains. **(A)** Schematic of unroofing and labeling procedure. Cells are grown on a coated glass coverslip and then unroofed leaving behind membrane sheets. Proteins are non-specifically labeled at cysteine residues using a maleimide-conjugated fluorophore (STAR-580 maleimide) and lipids are labeled by the addition of a membrane-partitioning dye (STAR RED-PEG2000-DPPE). **(B)** Representative STED images of unroofed membranes labeled for proteins (magenta; STAR-580 maleimide) and lipids (green; STAR RED-PEG2000-DPPE), showing relatively homogeneous lipid distribution but more punctate protein distribution. **(C)** Zoomed-in view of the square region from panel B, showing images acquired using STED (top) or confocal (bottom). **(D)** Representative intensity profiles for proteins (magenta) and lipids (green) along a line, showing anti-correlation of the signals. **(E)** Schematic showing the interpretation of the non-homogeneous protein signal and the anti-correlation with the lipid signal, namely that there exist protein-rich regions at the plasma membrane that tend to exclude the lipid label. **(F)** Representative confocal images of fixed, intact cells labeled for proteins (magenta; STAR 580-maleimide) and lipids (green; STAR RED-PEG2000-DPPE), showing that the labeling is primarily at the plasma membrane. **(G)** Representative confocal (top) and TIRF (bottom) images of proteins and lipids at the basolateral membrane. The protein signal is unevenly distributed and coincides with regions of reduced lipid signal, as in the unroofed membranes. Proteins are labeled with STAR 580-maleimide and lipids are labeled with STAR RED-PEG2000-DPPE.

We first looked at protein and lipid distributions in plasma membrane sheets obtained by unroofing adherent cells (**Fig. 1A**). We used shear flow for unroofing, but similar results were obtained by sonication following established protocols ^19^ (**SI Appendix, Fig. S1**). **Fig. 1B** shows a representative stimulated emission depletion (STED) microscopy ^20^ image of an unroofed cell membrane labeled for proteins (magenta) using STAR-580 maleimide and for lipids (green) using STAR RED-PEG2000-DPPE ^21^. While the lipid signal appears relatively homogeneous (with some variations that will be discussed), the protein signal is concentrated at certain spots rather than being a diffuse sheet. When looking at the protein and lipid signals more closely (**Fig. 1C**), most of the bright protein spots tended to be in regions where the lipid signal was weaker. This anti-correlation between the two channels (**Fig. 1D**) is accentuated in the STED data but is also visible in the confocal images (**Fig. 1C**). Additional images from different biological replicates and labeling probes are given in **SI Appendix, Fig. S2**.

What causes this anti-correlation between the protein and lipid signals? If we observed regions that are bright in both signals (which we do in some areas, see **Fig. 1B**), the simplest explanation would be some sort of local membrane topological variation ^22^ or vesicles that remain attached to the unroofed membranes, that would result in both more membrane signal and more protein signal. But instead, we see “shadows” in the lipid signal that are *sometimes* bright in the protein channel. To exclude artifacts related to specific labels, we tested different fluorophores for the protein and lipid labels, lipid reagents with different chemical structures, and omitted the protein label entirely (**SI Appendix, Fig. S3**), with similar results. We also ensured that the lighter and darker regions in the lipid signal are not a result of local membrane topology using *xyz* sectioning (**SI Appendix, Fig. S4**) ^23^.

We interpret the anti-correlation of the protein and lipid signals (**Fig. 1D**) to be due to “protein-rich domains” where lipids are partially excluded (**Fig. 1E**), perhaps simply by volume occupancy (or by some other means such as lipid composition). This explanation accounts for why not all of the shadows in the lipid signal are bright in the protein channel, because some of these domains presumably have proteins that are not labeled with the cysteine-reactive dye. We point out that we are not the first to describe domains that fit our description here; others have noted similar domains in studies using electron microscopy, atomic force microscopy, and fluorescence microscopy ^11–16^, but have not discussed the anti-correlation between lipid and protein labels.

What is the nature of these protein-rich domains? Some domains appear “stringy” while others appear roughly circular and both are presumably diffraction limited. The stringy appearance of some domains naturally raises the question of cytoskeletal elements. To test this, we labeled the unroofed membranes with actin stains (**SI Appendix, Fig. S5**) and found that these domains are only very sparsely labeled with actin markers, while actin labeling in intact cells was as expected. Furthermore, the fact that the lipid signal is depleted in these regions suggests that the protein species is present at the membrane itself, as proteins in the cytosol would not be expected to cause local depletion of the lipid stain. We thus conclude that both the stringy and the circular domains likely correspond to integral or peripheral membrane proteins, and perhaps the stringy domains represent proteins that tether the cytoskeleton to the cell membrane.

Do these protein-rich domains exist in intact cells, or are they an artifact of unroofing? To test this, we needed reagents that primarily label proteins and lipids at the plasma membrane but not inside the cell. We found that labeling for 5-10 minutes in serum-free media using STAR 580-maleimide (to label proteins) and the recently reported probe STAR-RED-PEG2000-DPPE (to label lipids) ^21^, followed by chemical fixation using PFA and then mounting of the coverslips resulted in staining primarily at the plasma membrane of these cells (**Fig. 1F**). We then imaged the plasma membrane at the adhesion surface either using a confocal microscope or using a total internal reflection fluorescence (TIRF) microscope (**Fig. 1G**) to examine protein and lipid distributions. The protein signal and lipid signal were anti-correlated in the cells, consistent with our findings in the unroofed membranes and with other studies that have shown their existence in live cell experiments ^12^. Thus, the plasma membrane of living cells appears to be organized into protein-rich domains and protein-poor domains.

### Protein-rich domains persist in giant plasma membrane-derived vesicles

Are these protein-rich domains only present in intact cells, or can they be isolated? We wondered whether they persist in cytoskeleton-free giant plasma membrane-derived vesicles (GPMVs), which are derived from cells by treatment with N-ethylmaleimide (NEM) and calcium (**Fig. 2A, B**) ^24,25^. GPMVs were isolated from cells (**Fig. 2A, B**) and burst onto the same glass coverslips used for the cellular imaging experiments. The membrane sheets were labeled using a membrane-partitioning dye (Cellmask Orange). Protein labeling was attempted but was unsuccessful, presumably due to NEM-capping of free cysteines. However, we reasoned that we could simply look at the variation in the lipid signal to infer the presence of protein-rich domains, (**Fig. 2C**), as we just concluded that in cells the lipid signal is anti-correlated with the protein signal (**Fig. 1D, E**). These experiments were carried out at ∼20°C, well above the temperature at which macroscopic phase separation occurs in these types of GPMVs ^25^. We note that since we are unable to directly observe the proteins in this experiment, inhomogeneous dye partitioning, local curvature differences, or bursting/surface-adhesion artifacts could contribute to lipid signal variation in GPMVs.

**Figure 2.**
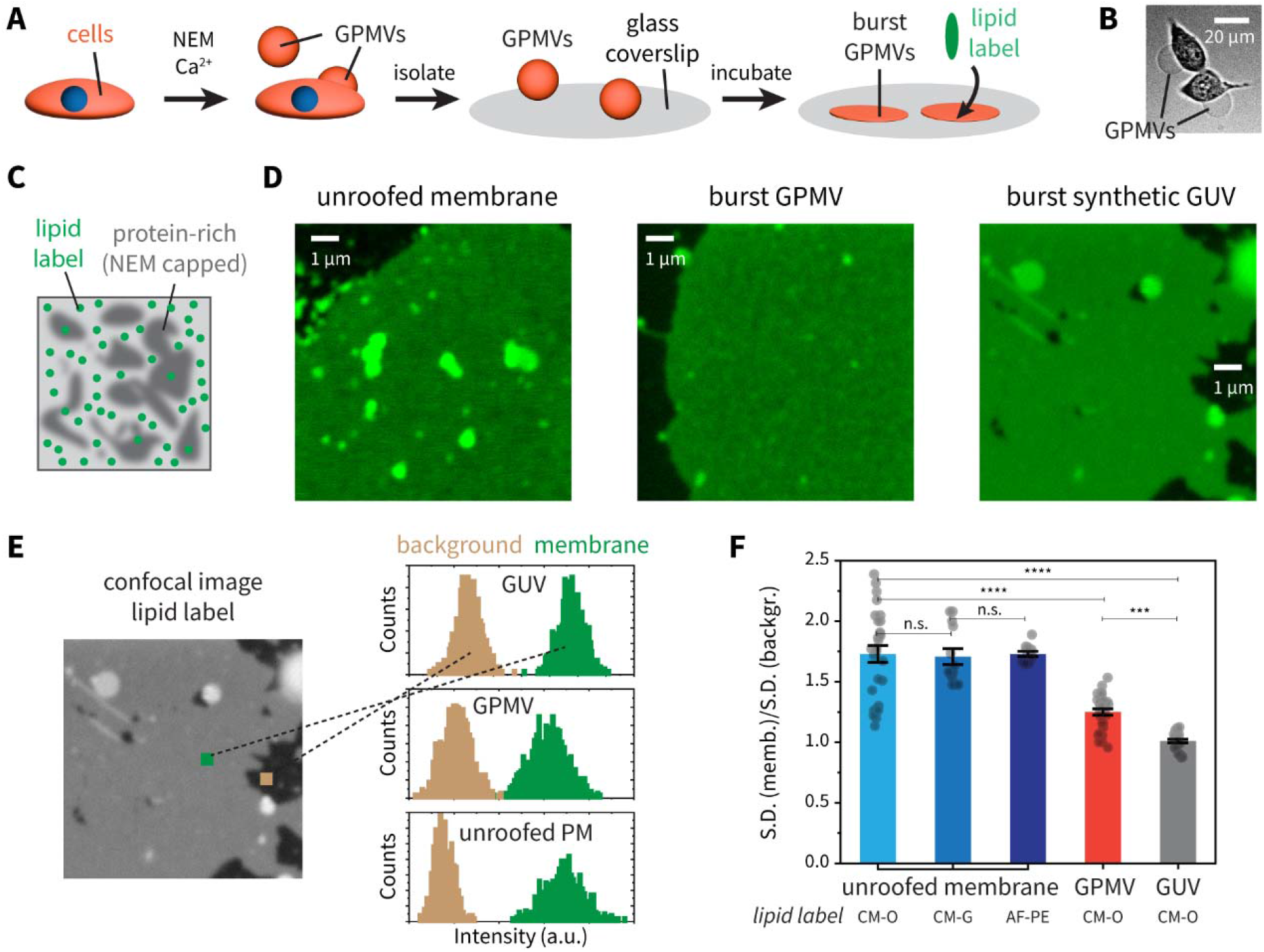
Protein-rich domains are also present in giant plasma membrane-derived vesicles. **(A)** Schematic of giant plasma membrane-derived vesicle (GPMV) generation from cells and their subsequent bursting for imaging. **(B)** Brightfield image of GPMVs budding off from cells. **(C)** Schematic showing the burst GPMV labeling method. As free cysteines are capped during NEM treatment, only the lipid label can be used and thus the protein-rich domains cannot be directly imaged using fluorescence microscopy. However, the presence of these domains can be inferred from the inhomogeneity of the lipid signal, which does not label the protein-rich domains as well as it does the protein-poor domains. **(D)** Representative confocal images of unroofed membranes from cells (left), burst cell-derived GPMVs (middle), and burst synthetic GUVs (right), all labeled using the same lipid partitioning label. **(E)** Principle of signal heterogeneity analysis in the membrane sheets: the width of the distribution of intensities in the background is compared to the corresponding distribution inside the membrane sheet. If the distributions are of similar width, then the signal heterogeneity can be explained simply by noise. **(F)** Plot of the ratio of the standard deviations of the distribution inside the membrane over that in the background (outside the membrane), a measure of the signal heterogeneity inside the membrane. If the ratio equals 1 (as is the case for GUVs), then the signal from the lipid label is taken to be homogeneous within the membrane. The data for unroofed cell membranes using three different lipid labels (Cellmask Orange, Cellmask Green, and AF594-PE) are shown. Cellmask Orange was used for the experiments with GPMVs and GUVs. The box represents the mean for each sample and the error bar is the standard error of the mean. The statistical test used was Welch’s T-test.

**Fig. 2D** shows representative results of lipid staining in unroofed plasma membranes, in burst GPMVs, and in lipid-only GUV controls. Variation in lipid signal intensity can clearly be seen in the unroofed membranes and in the burst GPMVs but are not visible in the GUV samples. To ensure that this phenomenon is not simply an optical illusion, we measured the distribution of intensities in small regions (chosen to not have any gross bright or dark spots) taken from either the membrane region or from the background in these images (**Fig. 2E**). The idea is that there is some natural variation in intensities due to noise. In the case of GUVs, the width of the distribution of intensities (a measure of the range of intensities in that region) is similar in the membrane sheet and in the background (**Fig. 2F**). Meanwhile in unroofed membranes and in GPMVs the signal from the membrane is noticeably broader than the background (**Fig. 2F**), but GPMVs show reduced variation compared to unroofed membranes. With the caveat that in this experiment we do not directly image proteins, we conclude that protein-rich domains likely persist to some extent in GPMVs. GPMVs are known to be free of cytoskeletal elements ^24,26^. Thus protein-rich domains likely arise spontaneously from the interactions within the mixture of proteins and lipids in the cell membrane.

### Purification of protein-rich and protein-poor vesicles

The data presented above, aligning with other studies ^11,12^, suggest that the cell membrane is organized into protein-rich domains. The data presented in this paper also indicate that protein-rich domains persist to some degree when isolating giant vesicles from cells. This finding suggests a way to isolate and analyze protein-rich and protein-poor regions from the cell membrane. After washing cells to remove secreted vesicles in the media, we generated GPMVs as described in methods and then gently sonicated them to produce smaller (sub-micron sized) vesicles we call small plasma membrane vesicles (SPMVs) (**Fig. 3A**). The size of SPMVs (∼50-100 nm in diameter) is around the size of protein-rich domains observed in cell membranes (∼100 nm or smaller). Cryo-electron microscope (cryo-EM) images show that some of the vesicles are filled with proteins and others are relatively empty (**Fig. 3B**). If proteins were randomly distributed in the membrane with, for instance, a Poisson distributed protein density, then the vesicles derived from the membrane should not show many nearly empty vesicles co-existing with many protein-packed vesicles. Why? Because for a random distribution of proteins, the presence of many protein-packed vesicles would predict very rare empty vesicles. By contrast, a cell membrane with protein-rich and protein-poor regions would be expected to generate vesicles with a multimodal distribution of protein density, as we see in cryo-EM micrographs.

**Figure 3.**
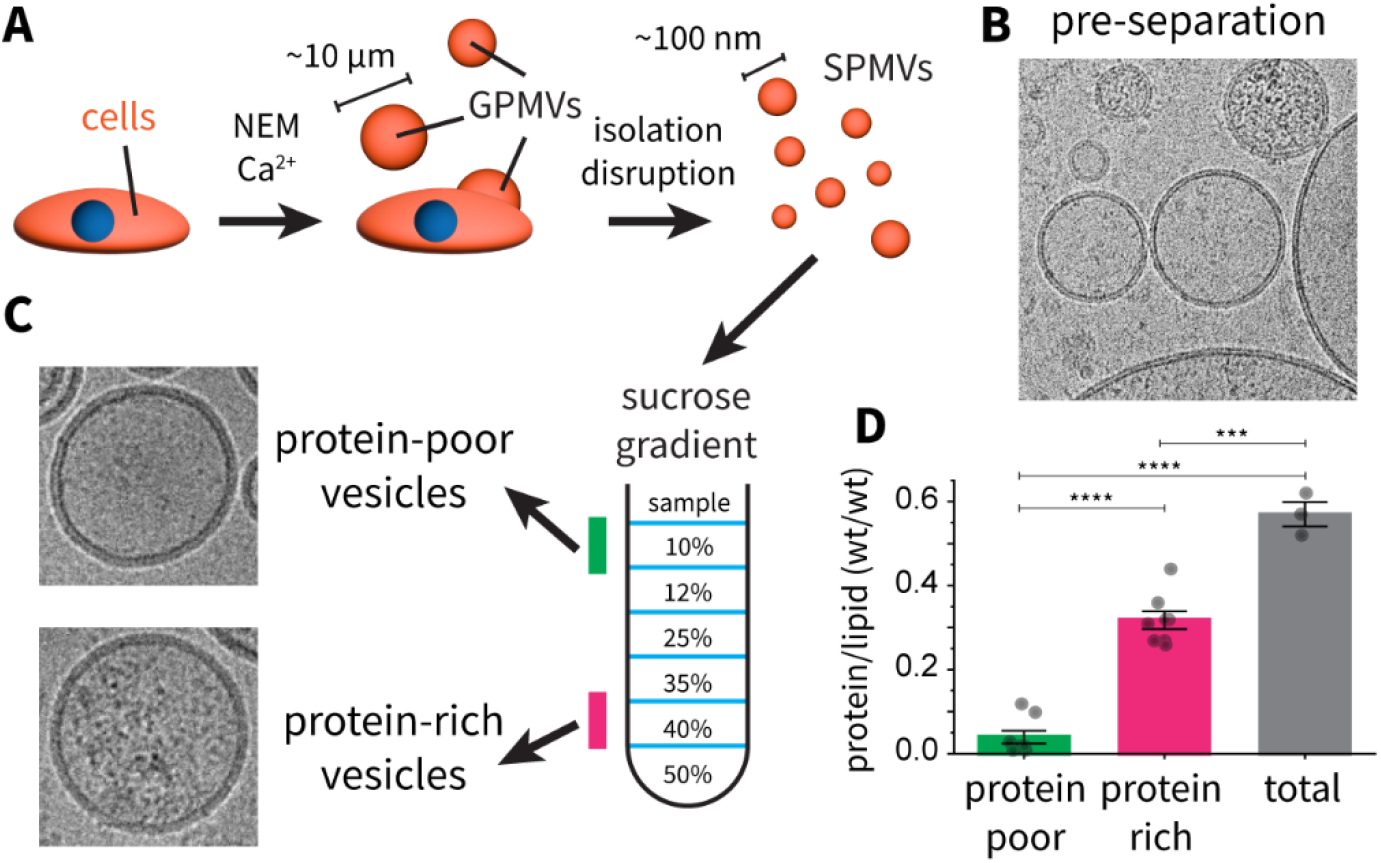
Isolation of protein-rich and protein-poor vesicles. **(A)** Schematic of small plasma membrane-derived vesicle (GPMV) generation from cells and their subsequent separation using a sucrose density gradient. **(B)** Cryo-EM image of cell-derived vesicles before separation, showing the presence of both vesicles with lots of protein and vesicles with little protein in them. **(C)** Representative cryo-EM images of protein-poor (top) and protein-rich (bottom) vesicles after separation using a sucrose density gradient. **(D)** Protein-to-lipid ratios (wt/wt), of the protein-poor, protein-rich and unseparated samples. The unseparated sample has a greater protein-to-lipid ratio because the bottom, most dense fractions from the sucrose gradient are not used. Protein concentration was calculated using a BCA assay and lipid concentration using ^1^H NMR. The box represents the mean for each sample and the error bar is the standard error of the mean. The statistical test used was Welch’s T-test.

The critical reader would be correct to wonder whether the protein-rich vesicles (**Fig. 3B, C**) are rich in membrane proteins or cytoplasmic proteins. We show in **SI Appendix, Fig. S6A**, that membrane proteins can have the appearance of being inside the vesicles rather than on the surface. But inspection of the membrane on the edge of vesicles shows that protein-rich vesicles contain many membrane proteins. Furthermore, we analyze (below) the protein composition of vesicles by mass spectrometry and find they contain membrane proteins and soluble proteins that are apparently associated with the membrane proteins. Consistent with this association, the abundance of membrane proteins versus total proteins is the same in the protein-rich and protein-poor vesicles (**SI Appendix, Table S1**). The important point is, the protein-rich vesicles are dense in membrane proteins.

To fractionate SPMVs into protein-rich and protein-poor vesicles we used sucrose density gradient ultracentrifugation (**Fig. 3C**) ^27^. There is, of course, a continuum of vesicle densities, but we chose fractions near the two extremes as representative examples (see **SI Appendix, Fig. S6B** for representative example of a sucrose gradient). Cryo-EM imaging of the purified vesicles confirmed that the different fractions from the sucrose gradient were indeed protein-rich or protein-poor (**SI Appendix, Fig. S6C**). To quantify the amount of proteins and lipids, we used a bicinchoninic acid (BCA) assay ^28^ to measure total protein concentration in the different samples and ^1^H nuclear magnetic resonance (NMR) spectroscopy ^29^ to quantify the total lipid concentration. The calculated protein-to-lipid ratios for the unpurified sample and the isolated protein-rich vesicles and protein-poor vesicles are shown in **Fig. 3D**. The protein-rich vesicles have about ten-fold more protein per lipid surface area compared to the protein-poor vesicles, and the ratio of protein and lipid in the protein-rich and unpurified samples is similar to published values on the composition of the plasma membrane ^30–32^, i.e., that proteins and lipids are about equal in abundance. The total amount of lipids is similar in the two fractions; thus, the protein-rich and protein-poor domains occupy roughly similar areas on the plasma membrane.

The above observations are consistent with a hypothesis that protein-rich and protein-poor vesicles are derived from protein-rich and protein-poor domains on the cell membrane (**Fig. 4A**). We next analyze the composition of these different vesicles.

**Figure 4.**
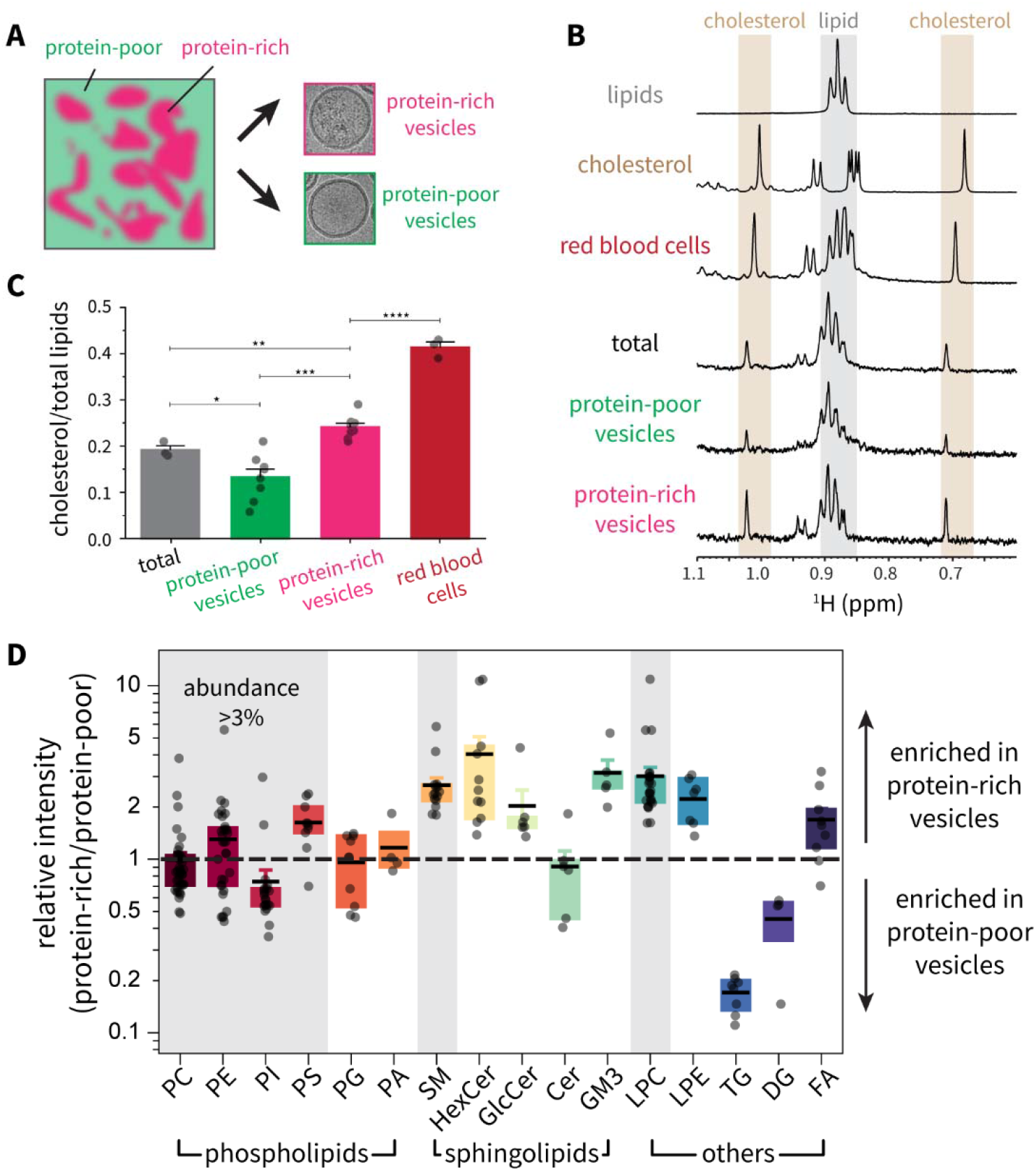
Lipid and cholesterol content of protein-rich and protein-poor domains. **(A)** Schematic of protein-rich domains and protein-poor domains at the plasma membrane, which are isolated as protein-rich and protein-poor vesicles. **(B)** Determination of lipid and cholesterol content using ^1^H NMR. The methyl region of the spectrum is shown with two resolved methyl signals from cholesterol and the methyl triplet from the terminal carbon in lipids. The top two spectra are from pure phospholipids and from cholesterol, respectively. **(C)** Cholesterol-to-lipid ratios (mol/mol) of the protein-poor, protein-rich and unseparated samples, and that of red blood cells determined using the same method as a control. The box represents the mean for each sample and the error bar is the standard error of the mean. The statistical test used was Welch’s T-test. **(D)** Relative intensities of different lipid species in the protein-rich vesicles divided by their intensities in the protein-poor vesicles, from lipid mass spectrometry. Lipid species are organized by headgroup, and each dot represents a unique lipid (with the indicated headgroup and a unique combination of fatty acid tails). Headgroups with the highest abundance according to mass spec intensity are marked with the grey shaded boxes. The box represents the middle 50% of the data (25-75% percentile range) and the error bar is the standard error of the mean.

### Cholesterol and ordered lipids are enriched in protein-rich domains

Lipid heterogeneity is thought to be important for plasma membrane organization *in vivo*. We thus measured lipid compositions in the two sets of vesicles. We measured cholesterol and total lipid content using ^1^H NMR as it is highly quantitative. The methyl region of the spectrum (**Fig. 4B**) shows the triplet from the terminal methyl of fatty acid chains in lipids, as well as two resolved methyl signals from cholesterol. We used the ratio of these lipid and cholesterol signals to calculate the fraction of cholesterol (cholesterol/total lipids) in the protein-rich vesicles, protein-poor vesicles, and in unenriched SPMVs as approximately 25%, 13%, and 20%, respectively (**Fig. 4C**). The cholesterol content in protein-rich vesicles is thus about two-fold higher than that in protein-poor vesicles. We performed similar recordings on red blood cells as a test of the accuracy our method. We found a cholesterol fraction of about 40% (**Fig. 4B, C**), in line with established reports on red blood cells ^33^.

To quantify other lipid species we used mass spectrometry. The relative enrichment of discrete lipid species (distinguished by their acyl chains), grouped by headgroup, in the protein-rich and protein-poor vesicles is shown in **Fig. 4D**. Lipids that are more ordered, such as sphingomyelin (SM), hexosylceramides (HexCer), and gangliosides (GM3), as well as phosphatidylserine (PS) are enriched in the protein-rich vesicles, whereas phosphatidylcholine (PC) and phosphatidylinositols (PI) are more abundant in the protein-poor vesicles, and phosphatidylethanolamine (PE) is equally distributed. We also analyzed enrichment across chain length and unsaturation (**SI Appendix, Fig. S7**) and found no clear trend amongst the different head groups. Cholesterol amounts were not measurable in these lipidomics experiment, but the NMR data addressed cholesterol. The enrichment of cholesterol in these domains raises the question whether depleting cholesterol disrupts these domains. Others have shown that cyclodextrin-mediated depletion of cholesterol not only disrupts these domains, but that the addition of exogenous cholesterol can promote the formation of these protein-rich domains ^12^. In summary, the protein-rich vesicles appear enriched in ordered lipids as well as cholesterol. A discussion of this finding in the context of the lipid raft hypothesis will follow later.

### Different proteins exhibit differential partitioning into protein-rich domains

If protein-rich vesicles originate from protein-rich domains, then we might expect there to be some differential segregation of certain proteins towards protein-rich domains and other proteins not. For example, some proteins might function in the crowded environment of a protein-rich domain, whereas others might function in a more dilute (protein-poor) environment. To address whether differential segregation exists, we performed bottom-up proteomics measurements on protein-rich and protein-poor vesicles normalized by the measured lipid and protein concentrations to account for the different protein density (per surface area) in the two fractions. Raw intensities for proteins were corrected within each sample using variance stabilizing normalization (VSN) ^34^. The normalized intensities were then corrected by the known protein and lipid concentrations (measured using the BCA assay and ^1^H NMR respectively; **Fig. 3D**) for each sample to adjust for the area of the cell membrane. The partition ratio (PR) for each protein was calculated as the ratio of the intensity of that protein in the protein-rich vesicles divided by the intensity of the same protein in the protein-poor vesicles (**Fig. 5A**), averaged over all pairwise sets of samples. For each protein with a PR, we mined the UniProt database ^35^ to obtain annotations for protein sub-cellular localization. Using this approach, we calculated partition ratios (PRs) for 4,258 proteins, of which about half are membrane proteins of different classes – multi-pass, single-pass, lipid-anchored and peripheral (**Fig. 5B**). The cytoplasmic proteins that were identified span broadly the cytoplasmic proteome.

**Figure 5.**
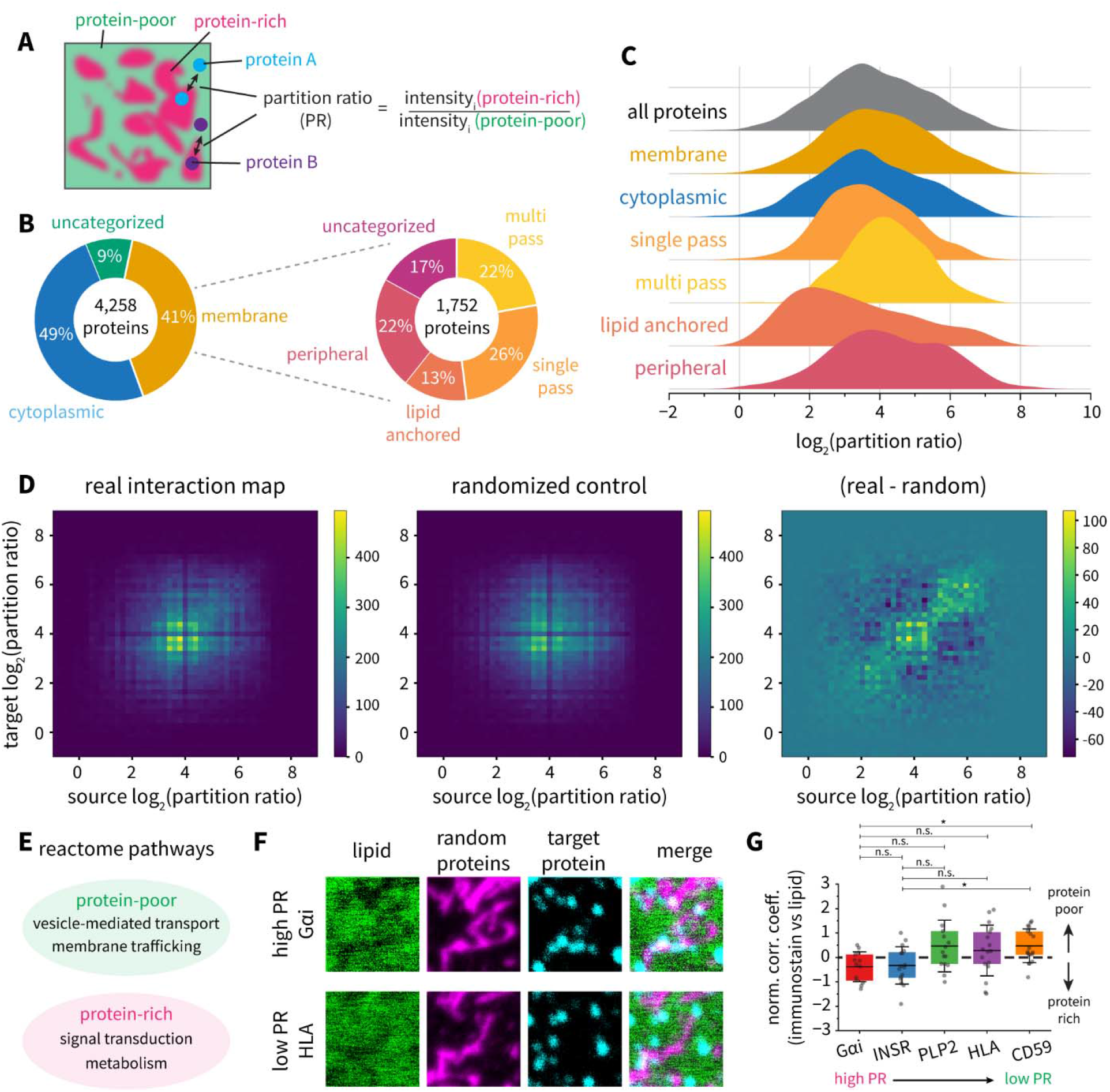
Protein content of protein-rich and protein-poor domains. **(A)** Schematic of the concept of protein partitioning between protein-rich and protein-poor domains. The partition ratio (PR) is defined as the ratio of intensities of a given protein in the protein-rich and the protein-poor vesicles. (B) Summary of the protein mass spectrometry data, showing the number and types of proteins characterized. About half of the total proteins are membrane proteins. 9% were uncategorized as either membrane proteins or cytoplasmic proteins based on their Uniprot identifiers. The 17% of uncategorized membrane proteins were not annotated according to membrane protein type. (C) Distributions of the partition ratio for all proteins and different subsets of proteins in the dataset. (D) Interactions between proteins in the dataset. Pairs of proteins (source and target) known to interact with each other were identified using STRING, and their partition ratios are plotted. Heatmaps of the interaction map are shown for the real data (left), a randomized control (middle), and the difference between the real data and the randomized control (right). The difference heatmap is linear, indicating that a subset of the proteins in the dataset (approximately a third) are known to interact with other proteins of similar partition ratios. (E) Reactome pathways enriched in the protein-poor and protein-rich domains, obtained from the subsets of proteins with the lowest (protein-poor) and highest (protein-rich) PR values, respectively. (F) Validation of the protein dataset by immunostaining proteins with different PR values in unroofed cell membranes. Representative images (2 µm x 2 µm) of immunostaining against a protein with high PR (Gαi, top row) and a protein with low PR (HLA, bottom row). Lipid signal is shown in green, the non-specific protein label (Cys-reactive) is shown in magenta, and the immunolabel against the specific protein is shown in cyan. (G) Summary of the immunolabeling data for five proteins, ranked by their PR values. For each protein, the normalized correlation coefficient between the lipid signal (which is depleted in protein-rich domains) and the specific immunolabel is shown. The normalization involved dividing by the absolute value of the correlation coefficient between the lipid and the protein signal, which was -0.1889 on average. The two proteins with higher PR have an anti-correlation coefficient that is about half of the lipid-protein anticorrelation in strength, while the proteins with lower PR show a weak positive correlation with the lipid signal. The box represents the middle 50% of the data (25-75% percentile range) and the error bar is the standard deviation. The statistical test used was Welch’s T-test, corrected for multiple tests using the Holm method.

The distribution of partition ratios for different subsets of protein types is given in **Fig. 5C**. Individual proteins are enriched to different extents in the protein-rich domains – some proteins are favored more than others. We point out that there are at least two reasons why a particular protein might have a high PR value: 1) the protein partitions favorably into protein-rich domains, or 2) the protein associates with another protein that partitions strongly into protein-rich domains. The PR distributions for most classes of proteins are similar — covering the whole range of PR values — but multi-pass membrane proteins tend to have higher PR values while lipid-anchored proteins (both GPI-linked and lipid tail anchored) have lower PR values on average (**Fig. 5C**). We will return to this in the Discussion, when considering what mechanism underlies the formation of these protein-rich domains.

What role might protein sorting into these domains play? We wondered whether proteins with similar partition ratios were known to interact with each other. To test this possibility, we identified all known unique interacting pairs of proteins (from the set of 4,258 proteins) using STRING functional network analysis ^36^, and plotted a symmetrized (i.e., for each unique source and target, there are two points that are symmetric about the diagonal) heatmap of their partition ratios (**Fig. 5D** – left panel). 2,721 of the 4,258 proteins had at least one interaction partner using the criteria used. If all interactions occurred between proteins of similar PR, we would expect a diagonal line. As a control, we randomized the pairing between the interactors and plotted the corresponding symmetrized heatmap on the same scale (**Fig. 5D** – middle panel). To see if there is a correlation between PR and function, we subtracted the randomized data from the real interaction map (**Fig. 5D** – right panel). What emerges is a clear linear trend – in other words, proteins that have similar PR values have been documented to interact as part of functional networks in living cells. This linear trend is accentuated for membrane proteins compared to cytoplasmic proteins in the dataset (**SI Appendix, Fig. S8**). With the caveat that STRING interactions do not necessitate spatial co-localization in the membrane, we suggest that organization into protein-rich and protein-poor domains could permit the spatial segregation of biological functions within the plasma membrane. Proteins with the highest PR values are involved in biological pathways such as signal transduction and metabolism, while proteins with the lowest PR values participate in vesicle-mediated transport and membrane trafficking (**Fig. 5E; SI Appendix, Fig. S9**).

Finally, to validate the measured protein partition ratios, we performed immunofluorescence experiments on five membrane proteins spanning the range of PR values to map them back onto protein-rich and protein-poor domains at the plasma membrane. We labeled unroofed membranes using a lipid label, a cysteine-reactive indiscriminate protein label, and an antibody against one of the five targets (**Fig. 5F**). Representative images for one protein (Gαi) with a high PR and one protein with a lower PR (HLA) are shown in **Fig. 5F** (other examples in **SI Appendix, Fig. S10**). As we saw before, the indiscriminate protein labeling is anti-correlated with the lipid stain in both cases.

However, most of the Gαi signal appears at or near these lipid shadows, which correspond to the protein-rich domains, while the HLA signal appears randomly distributed across the membrane (**Fig. 5F**). We quantified this phenomenon for the five targets by calculating the correlation between the lipid stain and the immunostain for the protein (**Fig. 5G**). Due to the low signal-to-noise, we normalized the correlation coefficient by the average correlation coefficient between lipid and protein (-0.1889; **SI Appendix, Fig. S10A**) which we know is negative (**Fig. 1D-E**). The two proteins with higher PR have an anti-correlation coefficient that is about half of the lipid-protein anticorrelation in strength, while the proteins with lower PR show a weak positive correlation with the lipid signal. In summary, these immunostaining experiments corroborate the partition ratios from the proteomics data and the connection between our studies of unroofed membranes and the cell-derived vesicles.

## Discussion

In the first part of this study we find that proteins in the plasma membrane of HEK cells are not randomly distributed but instead seem to exist in protein-rich domains on the order of 100 nanometers diameter, sometimes larger, separated by protein-poor regions (**Fig. 6**). While past studies have documented the heterogeneous nature of cell membranes ^11–16^, none have isolated domains to characterize their protein and lipid composition. That we have isolated protein-rich and protein-poor domains in this study hinges on the hypothesis that protein-rich and protein-poor vesicles derive from protein-rich and protein-pore domains. The observation that membrane heterogeneity persists at least partially in GPMVs, which are the intermediate vesicles between the intact cell membrane and the SPMVs, lends support for this hypothesis (**Fig. 2, D-F**). The demonstration that high and low PR proteins are correlated to protein-rich and protein-poor domains, respectively, measured in unroofed membrane sheets, also supports the hypothesis (**Fig. 5, F**,**G**). In the discussion below we therefore use ‘vesicle’ (the objects we isolated) and ‘domain’ (the objects from which the vesicles come from) interchangeably, with the understanding that this remains a hypothesis. For instance, domains could conceivably reorganize during vesicle formation or downstream biochemical processing.

**Figure 6.**
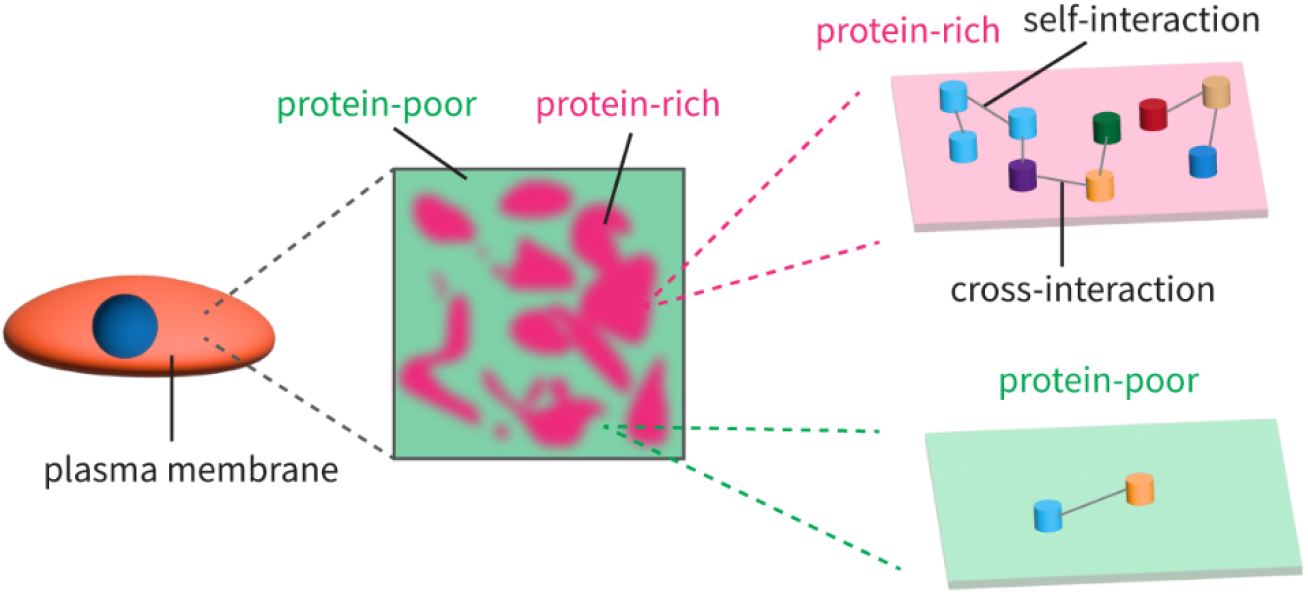
Schematic summarizing the protein-rich and protein-poor domains. The plasma membrane of living cells is organized into regions that are enriched in proteins, termed protein-rich domains, and regions that are poor in protein, called protein-poor domains.

Two major technical concerns tend to produce reflexive criticism of any attempts to interpret membrane organization. First, chemical fixation either due to incomplete immobilization or cross-linking effects can create artifacts. We note that others have demonstrated in live cells the existence of protein-rich domains similar to those we describe ^12^. Furthermore, while our data do not directly address the presence of these domains in living cells, they show that the domains are present in vesicles derived from cells that are not fixed. Second, the surface on which cells grow can influence membrane organization. But domains similar to those we describe have been observed in fixed cells grown on different substrates ^11^ as well as in live cells ^12^. There is no doubt that proteins in contact with a surface can be immobilized, and that surfaces can and do influence membrane organization, but this is reasonable – after all, free floating cells are scarce in our bodies. Moreover, to invoke surface interactions as the underlying cause of the organization we observe would necessitate patterning on the surface to direct the organization.

### Do protein-rich domains exist in different types of living cells?

Protein-rich domains have been observed primarily in studies imaging the distributions of specific proteins in a variety of different cell types grown on different surfaces ^11–16^. When looking at the distribution of specific proteins on the cell membrane using negative stain transmission electron microscopy, one finds that antibody labels against the protein are predominantly found at high-contrast, dark-appearing regions of the cell membrane ^10,11,14,15,37^. These results have been supported by atomic force microscopy measurements ^16^ and fluorescence microscopy of these domains ^12,16,17^, including in living cells. The presence and ubiquity of these domains, at least in the cell membranes of higher forms of life, seems likely. The existence of these domains thus modifies the picture of dilute, freely diffusing proteins in a sea of lipids, first painted by Singer and Nicholson in the 1970s ^38^, and adds to models of plasma membrane compartmentalization introduced since then ^1^.

The dearth of studies discussing protein-rich domains and the anti-correlation we observed between the lipid and protein signal is surprising to us, but closer inspection of many other studies shows that these domains are likely present in work by others but have just been missed. The anti-correlation has been mentioned in at least one study before ^12^, but is also visible in data reported by others: for instance in studies of GPCR domains at the plasma membrane ^6^, and of B cell receptor clusters in live cells ^7^. Finally, aside from the imaging experiments, it has long been recognized that the plasma membrane can be biochemically isolated as multiple different subfractions using density centrifugation ^39^. In fact, it has been shown before that simply homogenizing plasma membrane fractions of a given density can give rise to fractions of higher and lower density, corresponding to protein-rich and protein-poor membranes compared to the input membranes ^40,41^. Our observations should thus be pertinent to most studies of how proteins are distributed at the cell membrane, both in imaging experiments ^42^ and in single-particle tracking studies ^43^.

### Red blood cells: a model system for plasma membrane organization?

Red blood cells have long been used as a model system for the plasma membrane because these cells lack internal membranes – thus removing the requirement to isolate the outer membrane from the more abundant inner membranes. Freeze fracture studies of intramembrane proteins in red blood cell membranes show homogeneously and densely-packed proteins that are prone to aggregation upon treatment with trypsin or chemical agents ^44^. In contrast to such studies of red blood cells, freeze fracture experiments on the somata of neurons ^45^ show much sparser and more clustered intramembrane particles that align more closely with our experiments. The cholesterol levels we measure in our cell-derived vesicles are much lower than in red blood cells (**Fig. 4C**). The ‘textbook’ number of ∼40% cholesterol in the plasma membrane derives largely from studies on red blood cells ^33^; immortalized cell lines have approximately 22% cholesterol in their plasma membranes ^46^. In summary, we think that red blood cells are quite specialized in their function and may not always be representative of most other cell types.

### Implications for membrane-dependent biological processes

We have recently shown that many membrane proteins appear to self-assemble into homotypic – and in some cases heterotypic – small clusters in the cell membrane called higher-order transient structures (HOTS) ^10,47^. HOTS are formed by weak yet specific protein-protein interactions, which promote self-assembly between monomers diffusing in the cell membrane, and may increase the efficiency of signal transduction by locally concentrating molecules that function together, for instance, in a signaling pathway ^47^. It is notable that HOTS are concentrated inside protein-rich domains ^10^, as illustrated in **Figure 6**. Furthermore, others have shown that different kinds of proteins are non-randomly distributed inside protein-rich domains ^12^. In the present study we find that functionally linked proteins are correlated to their PR, which would be expected if functionally linked proteins are localized spatially (**Fig. 5D**). Taken together, these findings make plausible the idea that protein rich domains in cell membranes represent one of multiple levels of molecular organization in cell membranes. The protein rich domains represent relatively larger collections of different proteins, which then spontaneously self-organize inside the protein rich domains through other mechanisms such as specific self-assembly that underlies HOTS formation. This idea, if correct, naturally raises the question, what drives the formation of the protein rich domains in the first place?

### What is the mechanism of assembly of these protein-rich domains?

Protein-protein interactions, lipid rafts, and cytoskeleton-mediated corrals have all been proposed to contribute to plasma membrane organization. Here we find that isolated plasma membrane-derived vesicles, lacking cytoskeletal organizations, contain protein-rich and protein-poor domains. Protein-rich domains thus seem to form spontaneously from the protein-lipid mixture in the cell membrane. The cytoskeleton does likely still contribute to controlling the formation and size of these domains in cell membranes – for instance, it has been shown that disruption of cytoskeletal elements leads to larger clusters of many proteins ^11,48^. Perhaps the cytoskeletal mesh acts as a diffusion barrier, not just to confine proteins and lipids in certain zones, but also to prevent larger self-assemblies of proteins.

We found that protein-rich vesicles are rich in cholesterol and sphingolipids, thus, correlating protein density to these ordered lipids. However, we are unsure which came first. Do coalescing protein clusters bring these ordered lipids with them, or do the lipids form domains into which the proteins partition? The original observations surrounding the lipid raft model – that some proteins are resistant to detergent extraction – could be explained by either mechanism ^49,50^. We favor the idea that most membrane proteins (except some types of lipid-anchored proteins that are more akin to lipids) are inherently lipophobic to some extent, in that membrane proteins tend to associate in membranes to exclude the solvent (lipids), and that perhaps this process produces some form of phase separation ^51^. Analogous to protein folding in water ^52^, perhaps this process is driven by the entropy gained from lipids released from the annular shell upon protein association. The reason we favor a mechanism along the lines of this later picture is simply that most membrane proteins do not partition into liquid ordered (raft) phases when in the presence of liquid disordered (non-raft) phases (see **SI Appendix, Fig. S11** and ^53,54^ for examples).

It remains to be seen what precise mechanisms segregate the cell membrane into protein-rich and protein-poor domains. There must be identifiable molecular properties of proteins and/or lipids that drives this process. We deposit the lists of proteins and lipids enriched in protein-rich domains to promote further analysis to uncover the mechanism.

## Conclusion

In summary, the data here point to the existence of protein-rich and protein-poor domains in cell membranes and demonstrate how to isolate and characterize them. We conclude that most, but not all membrane proteins are concentrated in these protein-rich domains. Proteins that are functionally connected reside within these domains. Ordered lipids such as sphingomyelin and cholesterol are also enriched in these domains. Finally, the existence of these domains in the absence of cytoskeletal elements points to spontaneous assembly of components underlying their formation.

## Materials and Methods

### Cell culture

HEK293S GnTl^™^ cells were used in both adherent and suspension culture. Suspension cells to produce cell-derived vesicles were cultured in Freestyle 293 medium (GIBCO) supplemented with 2% fetal bovine serum, 100 U/mL penicillin, and 100 U/mL streptomycin at 37□ °C in 8% CO_2_. Adherent cells for imaging experiments were cultured in Dulbecco’s Modified Eagle Medium (GIBCO) supplemented with 10% fetal bovine serum and 2 mM L-glutamine.

### Preparation of intact and unroofed cells for imaging

Adherent HEK293S GnTl^™^ cells were grown either on high-performance glass coverslips (Zeiss, 0.17 mm thick) coated with poly-D-lysine (PDL) or on glass coverslips coated with both PDL and laminin (Corning BioCoat). For most experiments, cells were unroofed using a stream of Dulbecco’s phosphate-buffered saline (DPBS) through a 1 mL pipet applied at a ∼45° angle (∼10-15 applications per coverslip). Sonication (Qsonica Q125 sonicator equipped with a CL-18 probe) at a distance of ∼1 cm from the coverslip with 20% amplitude for ∼1 second was also used to ensure similar results as with shear flow.

Indiscriminate labeling of proteins was carried out by adding 15 µM of fluorophore-maleimide conjugate (eg. STAR-RED maleimide) in DMSO and incubating for ∼20 minutes at room temperature. Lipid labeling was accomplished by incorporating the Cellmask family of probes (Thermo Fisher) at dilutions of 1:1000 to 1:4000, or by adding 1µg/mL of AF594-PE (Avanti Research) in DMSO, or by adding 250 ng/mL of STAR-RED-PEG2000-DPPE (custom synthesis from Abberior) in DMSO. Typical lipid label incubation times were 5-20 minutes. Membranes were washed three times using DPBS to remove excess stain, fixed using 4% paraformaldehyde (PFA), washed again with DPBS three times, and then mounted on glass slides as detailed below.

The intact cell labeling was performed by incubating cells grown on the coated coverslips with STAR-580-maleimide (15 µM) and STAR-RED-PEG2000-DPPE (250 ng/mL) for 5-10 minutes at room temperature. Cells were then immediately washed and fixed using 4% PFA for 10 minutes, washed again, and then mounted on glass slides.

Coverslips with fixed, intact cells, or with fixed, unroofed membranes were mounted on glass slides with Prolong Diamond Antifade Mountant (Thermo Fisher), sealed using nail polish, and allowed to cure for 1-2 days before imaging.

### Preparation of GPMVs and GUVs for fluorescence imaging

Giant plasma membrane vesicles (GPMVs) were prepared from adherent cells grown in 6-well plates. Cells were washed with GPMV buffer (10 mM Hepes pH 7.4, 140 mM NaCl, 10 mM KCl, 2 mM CaCl_2_) twice before adding 2 mL of GPMV buffer with 7.5 mM N-ethylmaleimide (NEM)and incubating for 1.5 hours at 37°C and 5% CO_2_. 200 µL of the GPMV-containing solution was applied to a glass coverslip coated with PDL and laminin, let sit for 5 minutes, then GPMV buffer (without NEM) was added and the GPMVs were allowed to settle and burst for 1-2 hours. GPMVs were then labeled using Cellmask Orange (1:1000) for 20 minutes at room temperature, washed three times gently using NEM-free GPMV buffer, and then mounted on a glass side as detailed before.

Giant unilamellar vesicles (GUVs) comprised of synthetic lipids were prepared by dehydration-rehydration. Lipids (POPC or a 3:1:1 POPC:POPE:POPG mixture) were mixed in chloroform and dried to a thin film on 35 mm glass-bottomed dishes (Maktek) using a stream of argon and a vacuum dessicator for a few hours. The lipid film was rehydrated using 100 µL of GUV rehydration buffer (10 mM Hepes pH 7.4, 50 mM KCl, 200 mM sucrose) and let sit overnight at 4°C. The next day, the GUV-containing solution was harvested and applied to a PDL/laminin-coated coverslip, let sit for 5 minutes, and then GUV bath buffer (10 mM Hepes pH 7.4, ∼180 mM KCl, osmolarity matched) was added to the dish and the GUVs were allowed to settle and burst for 1-2 hours. GUVs were labeled by incubating with Cellmask Orange (1:1000) for 20 minutes at room temperature, washed three times with GUV bath buffer, and then mounted on a glass slide.

### Confocal/STED microscopy

Confocal and STED images were collected on a Facility line STED microscope (Abberior Instruments). The microscope was equipped with an Olympus IX83 stand, Olympus UPLXAPO ×100/1.45 NA oil objective, pulsed excitation lasers with time gating (405, 488, 561, and 640 nm) and a 775-nm pulsed STED depletion laser, 4 Avalance Photo Diode detectors, adaptive illumination packages (DyMIN/ RESCue), and a deformable mirror for correction of spherical aberrations. The Abberior Imspector software was used for image acquisition. Fluorophore excitation was performed at 488 nm (AF488-derivatives or Cellmask Green) 580 nm (AF594-derivates, STAR 580-derivatives or Cellmask Orange) and 640 nm (STAR RED-derivatives), whereas depletion for STAR-580 or STAR-RED was achieved using a wavelength of 775 nm. A pixel size of 15 nm was used in 2D STED mode. Excitation laser power, depletion laser power, line averaging/accumulation, and pixel dwell time were optimized for each imaging session to balance the signal-to-noise ratio and the resolution of the STED images. Images were exported as OBF files and visualized and analyzed using Fiji/ImageJ.

### TIRF microscopy

TIRF imaging was performed using a Nikon Eclipse Ti2-E inverted microscope equipped with a Tokai Hit stage-top incubator. Laser illumination was provided by a multimodal laser launch with wavelengths of 405 nm, 488 nm, 561 nm, and 647 nm. The laser beam was directed through a Nikon CFI SR HP Apochromat TIRF 100x oil-immersion objective (NA 1.49, WD 0.15 mm). The microscope was equipped with a Gataca iLas 2 controller for azimuthal TIRF control. The angle of illumination was adjusted to achieve total internal reflection, and the penetration depth of the evanescent wave was calibrated to ∼70-100 nm. Fluorescence emission was collected after passing through the appropriate filter and captured by a Hamamatsu ORCA-Fusion BT camera. Images were acquired sequentially for each channel with typical exposure times of ∼ 100 ms and laser power was set at about 20% of the maximum. Finally, sample focus was maintained using the Nikon Perfect Focus System and the system was controlled using Nikon NIS-Elements.

### Image visualization and analysis

Images were visualized using Fiji/ImageJ ^55^. The histogram analysis was performed by drawing a ∼300 nm-sided square and reading the standard deviation from the Histogram tool. Multiple locations within each image were taken for both the background and the membrane, and visibly brighter regions (assumed to be vesicles or incompletely unroofed regions) or holes in the membrane were excluded. Pearson cross-correlation coefficients between different channels were calculated using the inbuilt plugin, using multiple squares of ∼2µm x 2µm within each image. Again, notably brighter regions and holes were excluded as these would bias the calculation.

### Production and purification of protein-rich and protein-poor SPMVs

The protocol for production of SPMVs is very similar to that reported before ^18^. 1L of HEK293S GnTl^™^ cells were grown to a density of ∼3 × 10^6^/mL. Cells were harvested by pelleting at 500 × g for 10 minutes. The supernatant was decanted and the cells were gently resuspended in 200 mL of GPMV buffer to remove residual media and secreted vesicles (10 mM Hepes pH 7.4, 140 mM NaCl, 10 mM KCl, 2 mM CaCl_2_). The cells were harvested again at 500 × g for 10 minutes and resuspended in ∼400 mL of GPMV buffer with 7.5 mM NEM. Around 100 mL of this suspension was transferred to each of four 250 mL flasks, and incubated at 37°C and 8% CO2 for 1.5 hours with shaking at 130 rpm (25 mm throw; Infors HT). The flasks were shaken by hand to dislodge any attached vesicles, and the suspension was spun at 3,000 × g for 10 minutes to remove cell debris and large vesicles.

10% glycerol was added to the harvested vesicles and a probe sonicator (Branson 250, 1/2” tip) at 40% amplitude for three 30 s pulses with ∼1 min chilling on ice in between pulses. The vesicles were then pelleted by ultracentrifugation at 38,000 rpm (SW 70 Ti rotor) at 4°C for 40 minutes, and each pellet was resuspended by trituration in 1 mL of GPMV buffer with 10% glycerol and sonicated twice for 10 seconds in a bath sonicator (Branson M1800). Aggregates were removed by pelleting at 3,500 × g for 10 minutes and the vesicles were pooled and loaded onto sucrose density gradients (12%, 25%, 35%, 40%, and 50% wt/vol sucrose steps) prepared in ultracentrifuge tubes. Around 3 mL of sample was loaded on top of each gradient and spun for 20-24 hours at 38,000 rpm (∼170,000 × g; SW 41 Ti) and 4°C. Fractions of ∼1 mL in size were collected from the gradients after the spin and were individually washed (to remove sucrose) and concentrated in 2 mL 100 kDa MWCO spin filters (Amicon) until a final volume of ∼50 µL for each fraction. These fractions were aliquoted, flash frozen using liquid nitrogen, and stored at -80°C until use. Fraction number 1 and fraction number 6 from multiple experiments were used as the protein-poor and protein-rich fractions, respectively.

### Grid preparation and cryo-EM imaging

Grids for cryo-electron microscopy (cryo-EM) were prepared as described before ^56^. Quantifoil R1.2/1.3 400 mesh holey carbon Au grids were glow-discharged for 22 s, and 3.5 μL of concentrated membrane vesicles were applied and incubated for 3-5 minutes at 20°C with a humidity of 100%. The grids were blotted manually from the bottom edge of the grid with a piece of filter paper. Another 3.5 μL of sample was applied, and the grids were blotted using a Vitrobot Mark IV with a blot force of 0 and blot time of 3 s after 20 s of incubation. The grids were then flash-frozen in liquid ethane and stored in liquid nitrogen until data collection.

Cryo-EM images were collected on an FEI Talos Arctica transmission electron microscope operating at 200 kV. The microscope was equipped with an autoloader and a Gatan K2 camera. A target defocus of -3.5 µm was used to acquire the images.

### Protein and lipid quantification

Total protein content was quantified using a plate reader-based BCA assay (Thermo Fisher). The manufacturer’s protocol was used and 2% SDS was added to all the samples prior to the assay. Albumin was used to establish a linear calibration between 20 µg/mL and 2000 µg/mL. Protein content in the vesicle fractions was quantified using a plate reader (Tecan Infinite M1000) by measuring absorbance at 562 nm and using the calibration to calculate the protein concentration.

Lipid and cholesterol concentrations were measured using ^1^H solution NMR. A small amount of lipid vesicle fractions (∼10 µL) was dissolved in ∼550 µL of 2:1 (vol/vol) CD_3_OD:CDCl_3_ containing 100 µM TMSP as an internal standard and transferred to a 5 mm NMR tube for measurement. One dimensional ^1^H spectra were acquired using typically ∼32 transients on a 600 MHz (^1^H Larmor frequency) Bruker instrument equipped with an Avance Neo console and a 5mm HCN cryoprobe. Peak integration was carried out in Topspin after phasing and baseline correction and care was taken to avoid overlapping regions for quantification.

### Red blood cell lipid extraction

Extraction of total lipids from red blood cells (RBCs) was carried out using the Bligh-Dyer method ^57^. Red blood cells (Innovative Research) were washed three times with DPBS by pelleting at 3,000 × g for 5 minutes. Two preparations were tested that gave very similar results. First, washed RBCs were directly used for extraction by incubating 1 part of RBC slurry (∼80% water) with 2 parts of methanol and 1 part chloroform for 1 hour with agitation. Then 1 part of water and 1 part of chloroform were added, the mixture was filtered and allowed to phase separate and the bottom organic phase containing lipids was collected. The organic solvent was evaporated using a stream of argon gas and deuterated NMR solvent was added to the dried lipids to solubilize them. NMR measurements were carried out as detailed before. In the second method, RBC ghosts were first prepared by two rounds of hypo-osmotic shock (5 mM potassium phosphate buffer at pH 7.4, 0.2 mM EDTA, and 1 mM MgCl_2_) followed by four washes and pelleting ^58^, and then the lipids were extracted for NMR measurements as described for the first method.

### LC/MS lipidomics

LC-MS grade acetonitrile (ACN), isopropanol (IPA), methanol (MeOH), and 1-butanol (BuOH) were purchased from Fisher Scientific. High purity deionized water was filtered from Millipore (18 OMG). Lipid standards were purchased from Avanti Polar Lipids (Alabaster, AL, USA). Ammonium formate was obtained from Sigma-Aldrich in the best available grade.

Vesicles were extracted by adding 50:50 MeOH:BuOH + 10 mM ammonium formate at a 20:1 solvent-to-sample volume ratio. The vesicle–solvent mixture was subjected to bead-beating for 45 sec using a Tissuelyser cell disrupter (Qiagen), followed by 15 min sonication at 20°C. The mixture was centrifuged for 10 min at maximum speed. The above procedures were repeated twice to pool the supernatant. The final lipid extracts were dried in a Vacufuge (Eppendorf) and stored at -80ºC until analysis. On the day of lipidomics sample run, 40 µL of 50:50 MeOH:BuOH + 10 mM ammonium formate was added to the dry-down lipids, vortexed and sonicated to solubilize the lipids. The supernatant was transferred to sample vials for positive and negative ion lipidomics data acquisition.

The LC/MS platform for lipidomics analysis was as described previously ^59^, consisting of an Agilent Model 1290 Infinity II liquid chromatography system coupled to an Agilent 6550 iFunnel time-of-flight MS analyzer. An Agilent ZORBAX Eclipse Plus C18, 100 × 2.1 mm, 1.8 µm (Cat# 959758-902) reversed phase column was used for the separation. Mobile phases were (A) 10 mM ammonium formate with 5 μM Agilent deactivator additive (Cat# 5191-3940) in 5:3:2 water:ACN:IPA and (B) 10 mM ammonium formate in 1:9:90 water:ACN:IPA. Column temperature was set at 55°C and autosampler temperature was at 20°C. The flow rate was 0.4 mL/min with sample injection volume of 4 µL. The following gradient was applied: 0 min, 15% B; 0-2.5 min, to 50% B; 2.5-2.6 min, to 57% B, 2.6-9 min, to 70% B; 9-9.1 min, to 93% B; 9.1-11.1 min, to 96% B; 11.1-15 min, 100% B; 15-20 min, 15% B.

Raw data were analyzed using Mass Hunter Qualitative analysis (10.0), MassProfinder 10.0 and MassProfiler Professional (MPP) 15.1 software (Agilent technologies). Lipid peak chromatograms were extracted against an in-house database created using MassHunter PCDL manager 8.0 (Agilent Technologies) and lipid standards, based on monoisotopic mass (< 5 ppm mass accuracy) and chromatographic retention time matches (< 0.5 min). Matched lipids were further confirmed by comparing MS/MS fragmentation pattern to the corresponding lipid species standards.

Peak area ratios of extracted lipids were used to compare differences in partitioning into protein-rich and protein-poor vesicles, after normalizing each sample for the total integrated area of all identified lipids (to account for different amounts of lipids injected). The partition ratio for each lipid species was calculated as the pairwise ratio of the intensity of that lipid in the protein-rich vesicles divided by the intensity of the same species in the protein-poor vesicles.

### Protein mass spectrometry

SPMVs were diluted to afford solutions containing 0.8 µg of protein in 30 µL PBS (as measured by BCA), then sonicated for 5 minutes in a water bath. 3 µL of a stock of dithiothreitol (100 mM in water) was added to each sample, and the samples heated to 95 ºC for 5 minutes. 3 µL of a stock of iodoacetamide (200 mM in water) was then added to each sample and the mixtures incubated at room temperature in the dark. The SP3 protocol was then used to perform protein-level cleanup ^60^. Samples were then again sonicated for 5 minutes in a water bath. 7.5 µL of Sera-Mag SpeedBeads (a 1:1 ratio of GE cat. No. 45152105050250 and cat. No. 65152105050250, washed three times with water and suspended at a density of 50 mg/mL) was added to each sample, and 36 µL of acetonitrile (UHPLC-MS grade) added. These mixtures were then allowed to stand without agitation for 8 minutes at room temperature. A magnetic rack was used to pellet the beads, and the supernatant aspirated. Beads were then washed twice with 100 µL of UHPLC-MS grade 80% ethanol/water and twice with UHPLC-MS grade acetonitrile, each time with resuspension. Beads were then resuspended in 50 µL of ammonium bicarbonate in water containing 0.2 µg MS-grade trypsin (Pierce, cat. No. 90058, prepared from dilution from a 1 mg/mL stock solution in 50 mM acetic acid) and sonicated in a water bath for 30 seconds. The suspended beads were then incubated at 37 ºC for 16 hours with continuous inversion. Samples were then centrifuged (21,000G, 10 minutes) and the supernatant filtered using cellulose acetate cartridges (Corning Costar Spin-X Centrifuge Tube Filters, 0.22 µm pore CA membrane, PN 8161). The flowthrough was then desalted using Pierce C18 spin tips according to the manufacturer protocol with formic acid substituting trifluoroacetic acid, and dried peptide samples resuspended in 20 µL 2% UHPLC-MS grade acetonitrile/water containing 0.1% formic acid.

Data was obtained using a timsTOF Pro 2 mass spectrometer system integrated with a nanoElute UHPLC. A Waters nanoEase Symmetry C18 trap column (cat. No. 186008821, 5 µm particle size, 180 µm ID x 20 mm length) was used for online sample cleanup with an IonOpticks Aurora Ultimate column (75 µm ID, 1.5 µm particle size, 25 cm length) with integrated captivespray emitter used for analysis. A two-column separation method was used with a 60 minute gradient spanning 2-37% acetonitrile/water at a 150 nL/min flow rate, with the default long gradient DIA method used for MS data acquisition (1 second cycle time). Raw data was processed using DIANN using an in-silico spectral library generated from a human proteome to provide protein-level IDs using default settings ^61^.

Raw intensities obtained for each protein were corrected within each sample using VSN normalization in R ^34^. The normalized intensities were then corrected by the known protein and lipid concentrations for each sample to adjust for the area of the cell membrane. The partition ratio (PR) for each protein was calculated as the pairwise ratio of the intensity of that protein in the protein-rich vesicles divided by the intensity of the same protein in the protein-poor vesicles. We only report results for proteins that were detected in at least two independent samples for each set of vesicles. For each protein with a PR, we mined the UniProt database ^35^ to obtain annotations for protein sub-cellular localization and for basic structural information.

To query proteins known to interact, we used the STRING protein-protein interaction database ^36^ running on the Cytoscape StringApp ^62^. The 4,258 proteins that were identified from the mass spectrometry data were input into the STRING database and a cutoff of 0.8 was used to filter out less certain interactions. The output from STRING is a list of all unique (meaning each pair only shows up once, not twice) pairwise interactions between the proteins. 2,721 of the 4,258 proteins had at least one interactor that passed the cutoff filter. A Python script was used to obtain the PR values for all of the pairwise interactors identified from STRING, symmetrize the output (i.e. for each pair there are now two points that are symmetric about the diagonal), and then plot the symmetrized data as a heatmap with a defined number and width of bins and scales.

### Immunofluorescence experiments in unroofed membranes

Primary antibodies against each target were purchased and screened for non-specific labeling and optimal labeling dilutions before use. The following primary antibodies were used; Gαi: Thermo MA5-12800, INSR: Abcam ab245688, PLP2: Abcam ab208266, HLA-ABC: Thermo MA5-11723, and CD59: Thermo MA1-19133. AF488-labeled secondary antibodies against the appropriate host were used.

Cells were plated on PDL/laminin-coated coverslips. Residual media was removed by washing twice with DPBS. The cells were then unroofed by shear flow and then washed three times with DPBS. Indiscriminate protein and lipid labeling was carried out using STAR-RED maleimide and Cellmask Orange, respectively, as detailed before. The unroofed membranes were washed three times, blocked with 2% bovine serum albumin (BSA) in DPBS for 30 minutes, washed again two times, and then labeled with primary antibody for 45 minutes at the optimal dilution for each antibody (in the presence of 2% BSA). Excess primary antibody was removed by washing three times and the secondary antibody was then added at a dilution of 1:500 with 2% BSA for 45 minutes. Excess secondary antibody was removed by washing three times and the samples were fixed with 4% PFA, washed again, and then mounted on a glass slide for imaging.

### Preparation of uniform and phase-separated GUVs containing proteins

GIRK, Plp2, CD63, and Gβγ were expressed and purified as described before ^63–65^ and reconstituted into small unilamellar vesicles (SUVs) of the appropriate lipid composition (soy PC, 2:2:1 or 1:2:2 DOPC:egg SM:cholesterol, all from Avanti Polar Lipids) as detailed before ^66^. 0.1%-0.4% Rhodamine-PE (18:1) or Ganglioside GM1 (ovine brain) were doped in as fiducials for the liquid disordered and liquid ordered phases, respectively, as necessary. GUVs were generated by electroformation ^64,67^ using indium tin oxide glass slides in an incubator oven set at ∼50°C. GM1-labeling was performed by addition of AF488-cholera toxin B (Thermo) after GUVs were made. GUVs were imaged on 35mm collagen-coated glass-bottomed dishes (Maktek) using an ECHO Revolve microscope equipped with a 100x oil objective.

## Data Availability

Partition ratios from the proteomics and lipidomics analysis are deposited with the manuscript. All other data is available from the authors upon request.

## Supporting Information

SI Appendix: additional fluorescence microscopy data, functional network analysis, lipid MS analysis, and cryo-EM images.

## Acknowledgments

We thank Priyam Banerjee, Alison North, and other members of the Bio-Imaging Resource Center for their assistance with fluorescence imaging experiments. We thank Johanna Sotiris, Honkit Ng, and Mark Ebrahim at the Evelyn Gruss Lipper Cryo-EM Resource Center for assistance with cryo-EM grid screening. We thank Sean Brady for providing access to the NMR spectrometer, and Yi Chun Hsiung for assistance with insect and mammalian cell cultures. Finally, we thank members of the MacKinnon Lab for helpful discussions, and Dr. Jue Chen and her research group for their advice. V.S.M. is supported by the Jane Coffin Childs Memorial Fund Fellowship. R.M. is an investigator in the Howard Hughes Medical Institute.

## TOC/Graphical Abstract

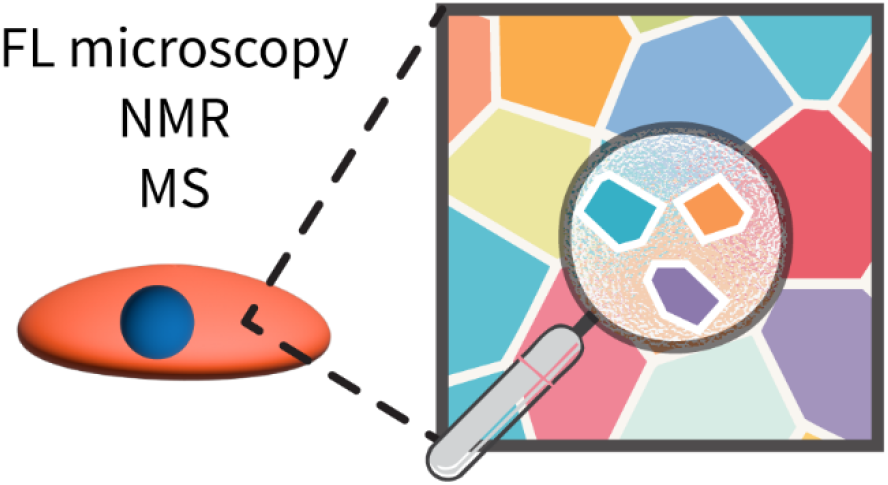

