## Supplementary material for "Spatial deconstruction of the plasma membrane": SI Appendix

^◆^ S.G. deceased on September 1, 2025.

**This PDF file includes:**

Figures S1 to S11

Table S1

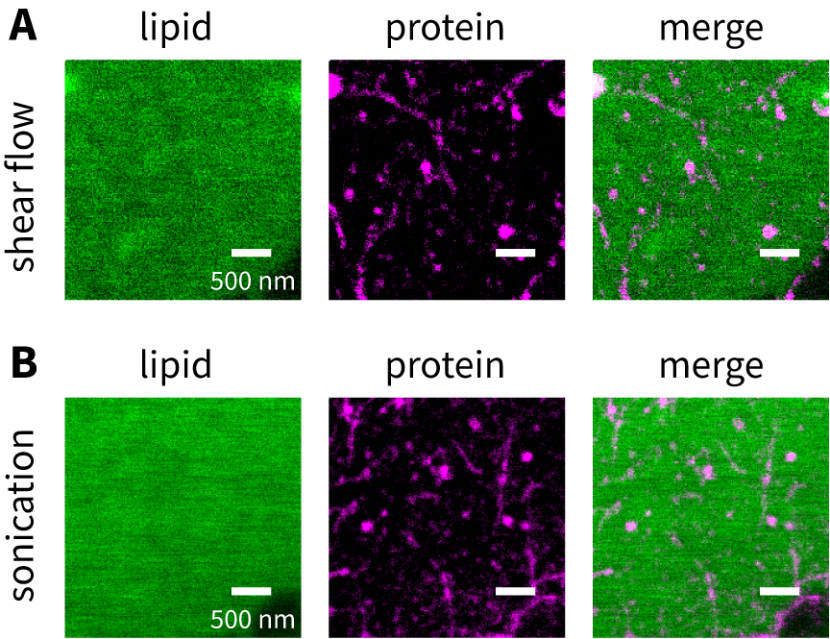

**Figure S1.** Unroofing by shear flow and sonication gives similar results.

Representative confocal (lipid) or STED (protein) images of unroofed membranes prepared by (**A**) shear flow, or (**B**) sonication. The unroofed membranes were labeled for proteins (magenta) and lipids (green).

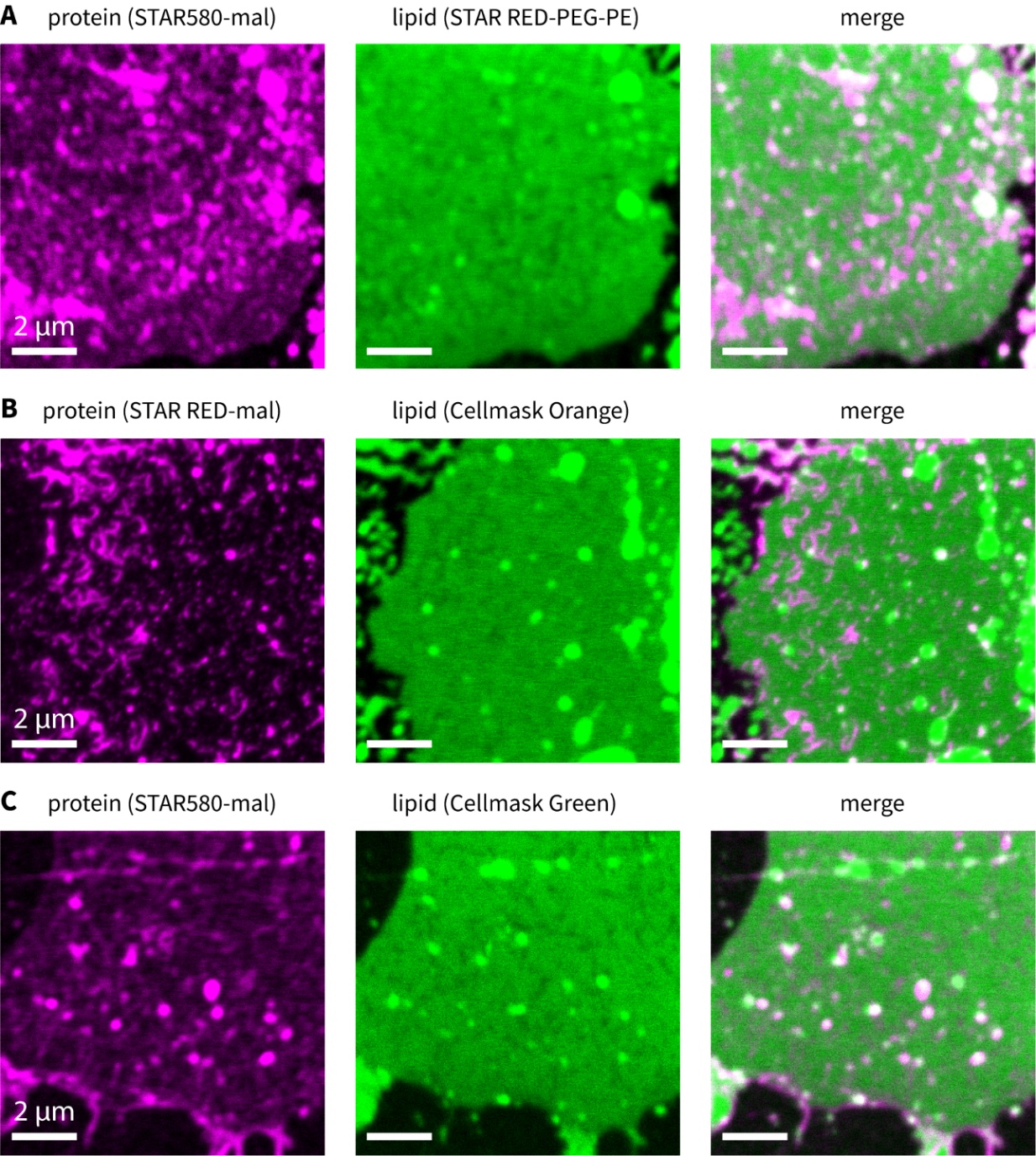

**Figure S2.** Additional images showing protein-rich regions in unroofed plasma membranes.

Representative confocal images of unroofed membranes labeled for proteins (magenta) and lipids (green) from different batches of biological replicates. Each image is the uncropped confocal image acquired and the three images were measured using different combinations of labels. (**A**) Protein labeled using STAR580-maleimide, lipid labeled using STAR RED-PEG-PE. (**B**) Protein labeled using STAR RED-maleimide, lipid labeled using Cellmask Orange. (**C**) Protein labeled using STAR580-maleimide, lipid labeled using Cellmask Green.

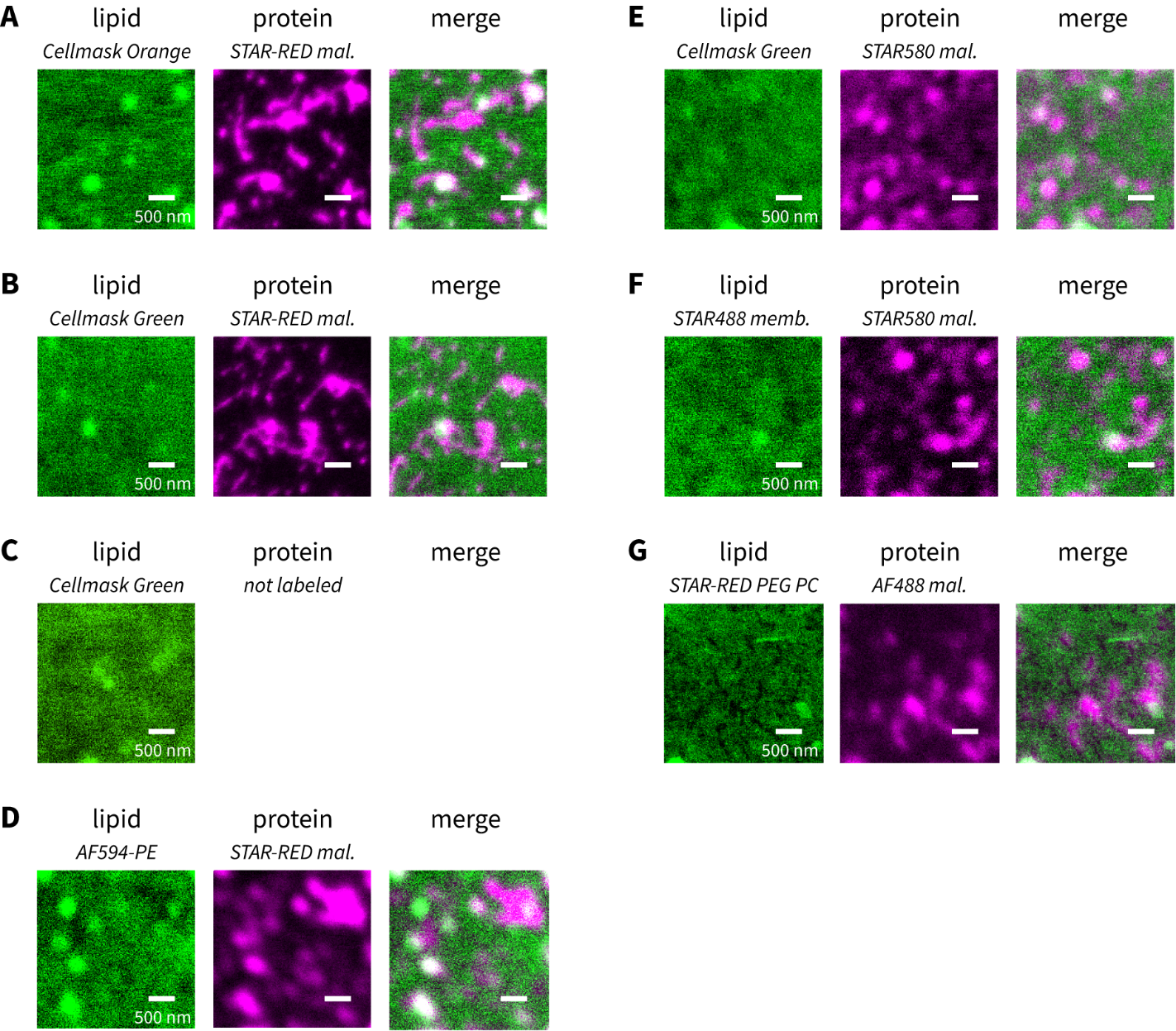

**Figure S3.** Anticorrelation between proteins and lipid signals is independent of label choice.

Representative images of unroofed membranes labeled for proteins (magenta) and lipids (green) using different combinations of fluorophores. (**A**) Lipids labeled using Cellmask Orange (ex. 556 nm) and proteins labeled using Abberior STAR-RED (ex. 638 nm) maleimide. (**B**) Lipids labeled using Cellmask Green (ex. 522 nm) and proteins labeled using Abberior STAR-RED (ex. 638 nm) maleimide. (**C**) Lipids labeled using Cellmask Green (ex. 522 nm), while the protein label was excluded. (**D**) The fluorescent lipid 18:1 PE-Topfluor AF594 (ex. 590 nm) was incorporated and proteins were labeled using Abberior STAR-RED (ex. 638 nm) maleimide. (**E**) Lipids labeled using Cellmask Green (ex. 522 nm) and proteins labeled using Abberior STAR-580 (ex. 587 nm) maleimide. (**F**) The fluorescent lipid STAR-488-membrane (ex. 503 nm) was incorporated and proteins were labeled using Abberior STAR-580 (ex. 638 nm) maleimide. (**G**) The fluorescent lipid STAR-RED-PEG-PE (ex. 638 nm) was incorporated and proteins were labeled using AF488 (ex. 490 nm) maleimide.

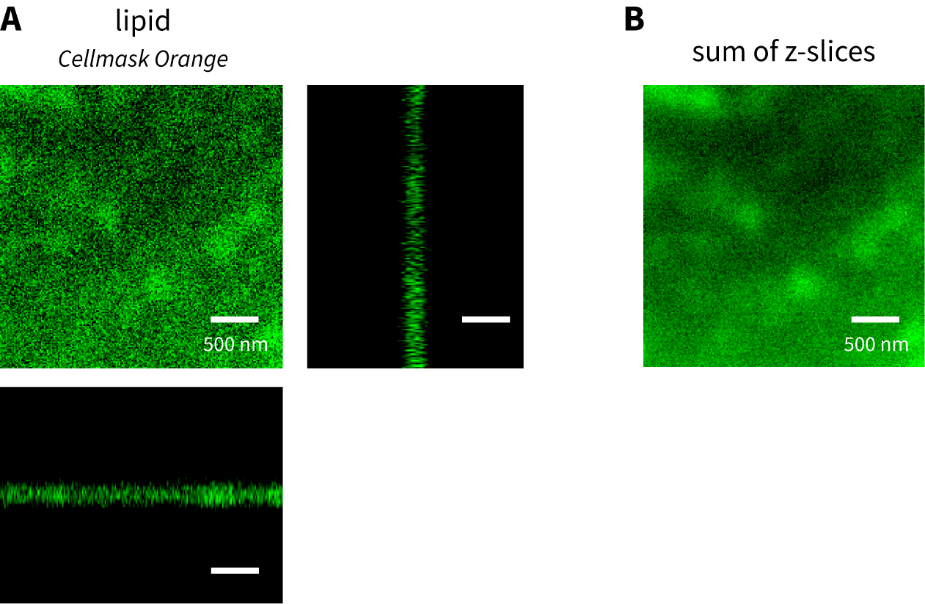

**Figure S4.** Variation in lipid signal intensity is not due to local curvature differences.

(**A**) Orthogonal slices of an unroofed membrane labeled with a lipid stain (Cellmask Orange). Some regions of the *xy* plane appear darker than others, not because these regions are out of focus compared to other regions. The three panels represent orthogonal views of the z-stack, showing that there is no detectable curvature in the *xz* and *yz* planes. (**B**) This can be confirmed by plotting the sum of z-slices, which shows the same intensity distribution as the single *z* slice at the unroofed membrane shown in panel A. Scale bar is 500 nm.

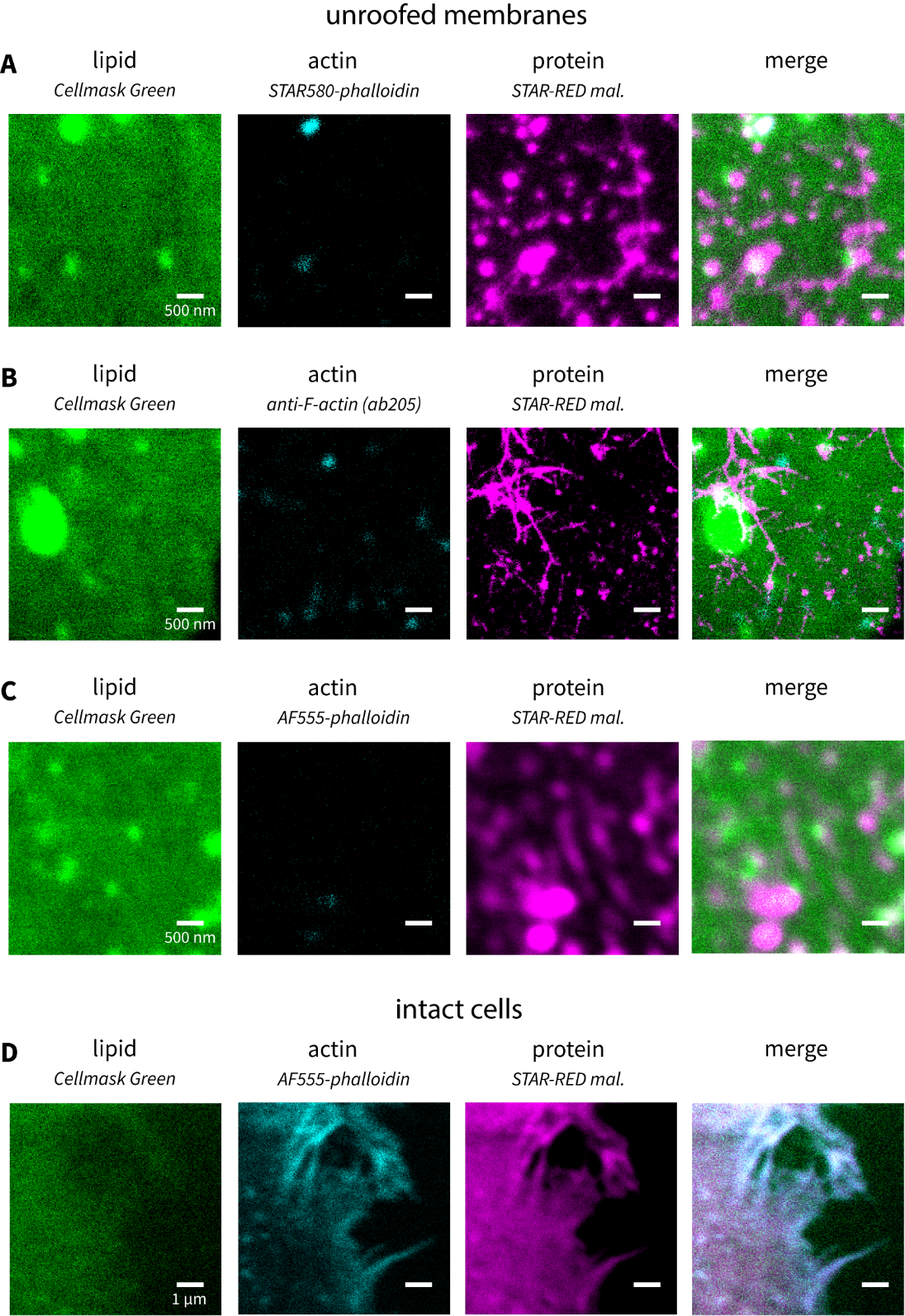

**Figure S5.** Actin labeling shows limited cytoskeleton retained in unroofed membranes.

Representative images showing lipid, actin and protein labeling in (**A-C**) unroofed membranes and in (**D**) intact cells. Lipids and proteins were labeled using Cellmask Green and STAR-RED maleimide in all panels. Actin was labeled using phalloidin conjugated to STAR-580 (**A**) or AF555 (**C** and **D**), or using an anti-F-actin antibody (ab205) further labeled with a STAR580-labeled secondary antibody (**B**). While actin staining in intact cells is clearly visible, staining in the unroofed membranes is very limited.

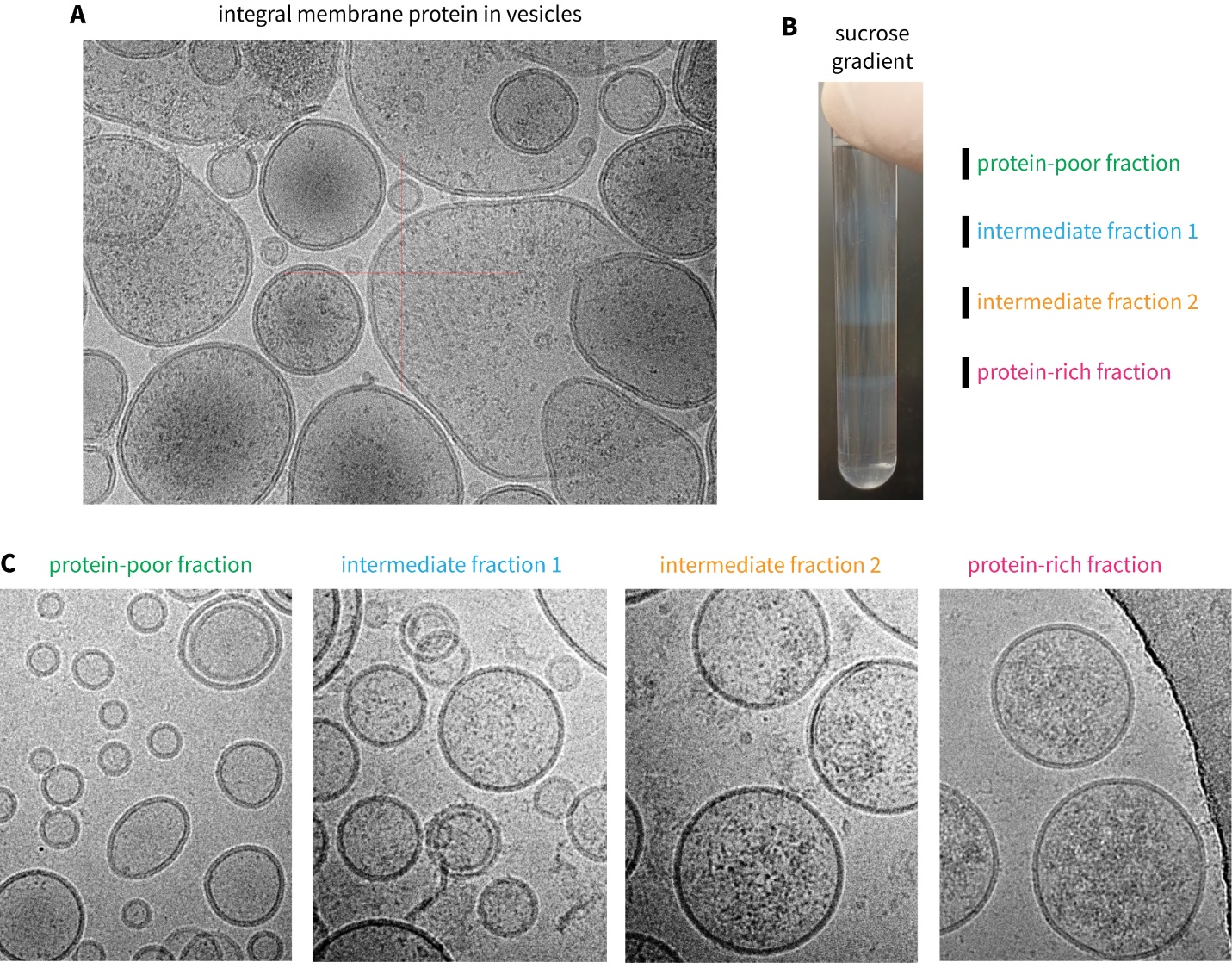

**Figure S6.** Cryo-EM images of isolated protein-poor and protein-rich vesicles, and an integral membrane protein reconstituted into vesicles for comparison.

(**A**) A representative cryo-EM micrograph of an integral membrane protein, Kv2.1, reconstituted into lipid vesicles. Notice how the membrane proteins appear to be “inside” the vesicle because the image is a projection and includes the top and bottom membranes. (**B**) Representative image of sucrose density gradient used for purification. Positions of fractions used for panel (**C**) are indicated. (**C**) Representative cryo-EM images of (left to right) the protein-poor fraction, two intermediate fractions, and the protein-rich fraction purified using the sucrose density gradient.

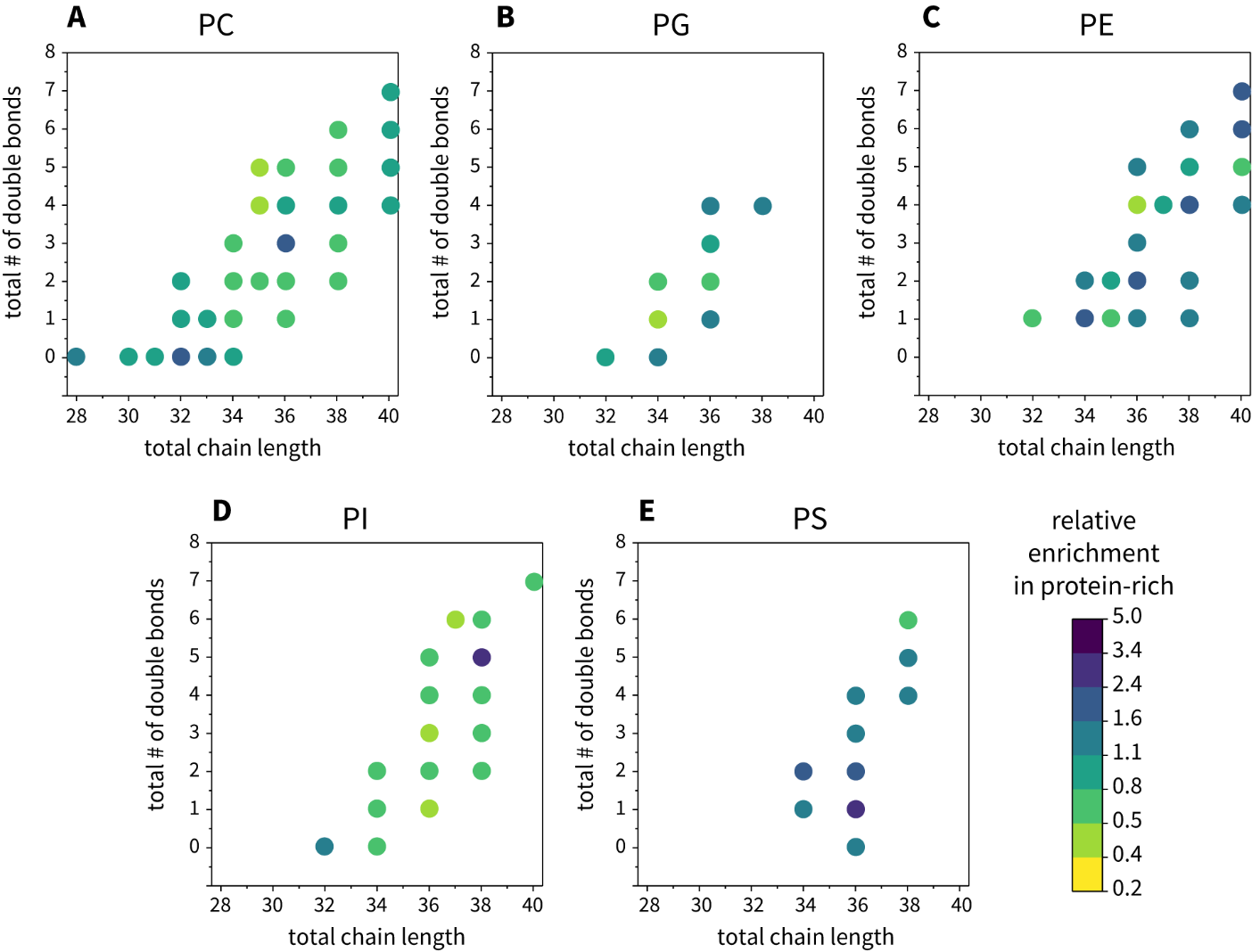

**Figure S7.** Lipid acyl chain length and unsaturation are not determinants of sorting into protein-rich domains.

Relative intensities of different lipid species in the protein-rich vesicles, plotted against the total chain length of the species and the total number of double bonds in the species. The species are sorted by headgroup: (**A**) PC, (**B**) PG, (**C**) PE, (**D**) PI, and (**E**) PS. There is no clear trend within each headgroup, but some headgroups are clearly more enriched in the protein-rich vesicles on average (for instance PS).

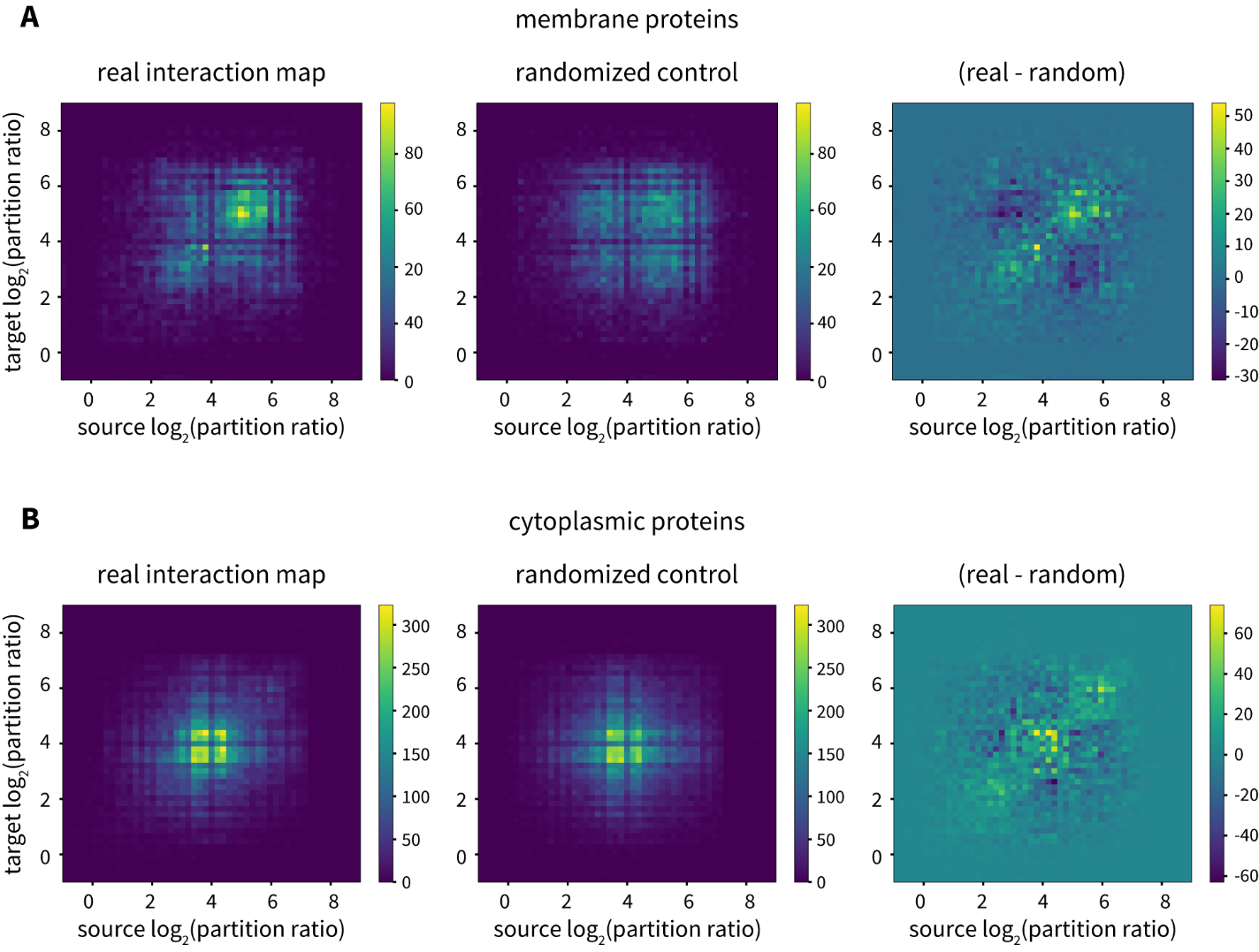

**Figure S8.** Interaction analysis for membrane proteins and for cytoplasmic proteins.

STRING-identified interactions between (**A**) membrane proteins and other proteins and (**B**) cytoplasmic proteins and other proteins in the dataset. Pairs of proteins (source and target) known to interact and their partition ratios are plotted. Heatmaps of the interaction map are shown for the real data (left), a randomized control (middle), and the difference between the real data and the randomized control (right).

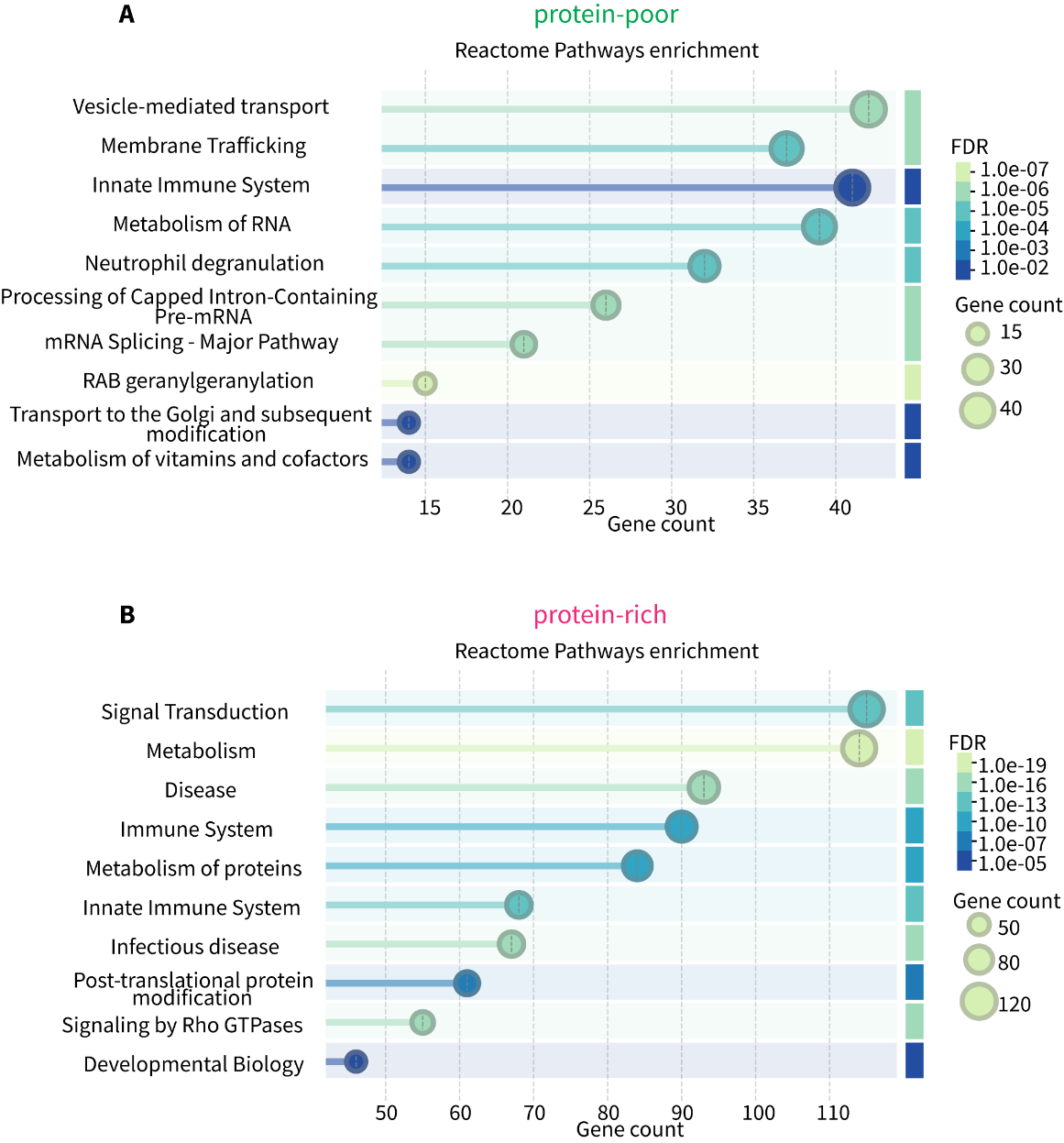

**Figure S9.** Reactome pathway analysis for high-PR and low-PR proteins.

(**A**) The 500 proteins with the lowest partition ratios (PR) and (**B**) the 500 proteins with the highest partition ratios were used to infer Reactome Pathways enriched in the protein-poor and protein-rich domains, respectively). The pathways are sorted by gene counts (reported in absolute counts, out of the input of 500 proteins). The key for the False Discovery Rate (FDR) is given on the right.

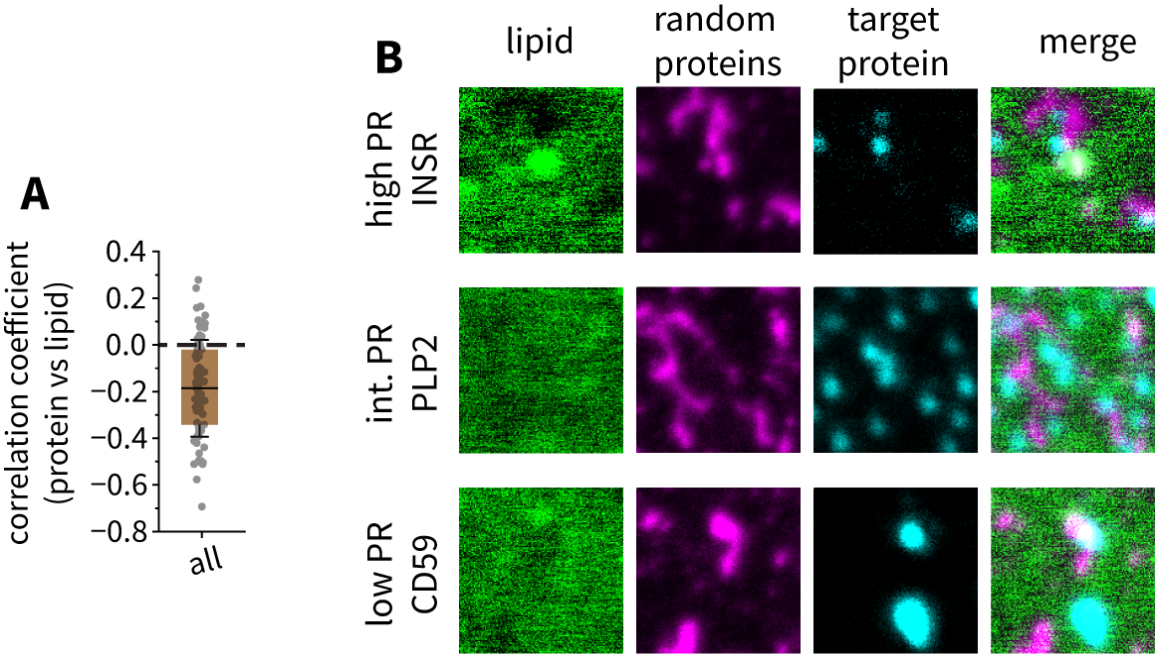

**Figure S10.** Additional data for immunostaining analysis of proteins with different PRs.

(**A**) Pearson correlation coefficient between the lipid signal (which is depleted in protein-rich domains) and randomly labeled proteins (using the cysteine-reactive dye). The data are averaged across all the proteins tested for immunolabeling because results were similar for the different proteins. (**B**) Additional images (2 µm x 2 µm) of immunostaining against proteins with high PR (INSR, top row), intermediate PR (PLP2, middle row), and low PR (CD59, bottom row). Lipid signal is shown in green, the non-specific protein label (Cys-reactive) is shown in magenta, and the immunolabel against the specific protein is shown in cyan.

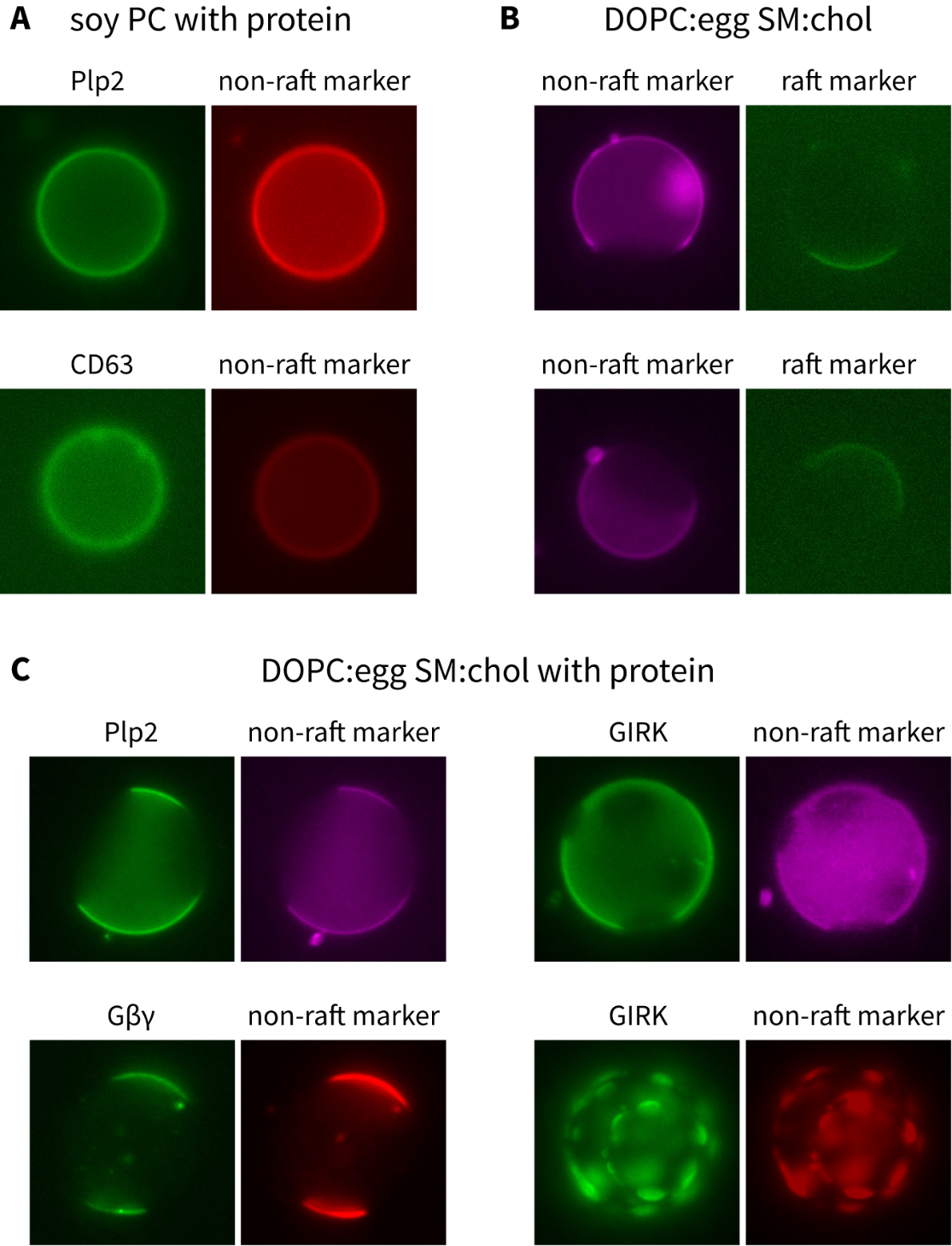

**Figure S11.** Partitioning of proteins between non-raft and raft domains in synthetic GUVs.

Representative epifluorescence images of GFP-labeled proteins and fluorescent lipid labels in uniform and phase-separated GUVs. (**A**) Two integral membrane proteins (Plp2 and CD63, both GFP-tagged) in soy PC GUVs labeled with rhodamine-PE. In non phase-separated GUVs the proteins are uniformly distributed. (**B**) Raft (AF488-cholera toxin) and non-raft (rhodamine-PE) labeling in phase separated GUVs consisting of a mixture of DOPC, egg sphingomyelin, and cholesterol. The liquid ordered (raft) and liquid disordered (non-raft) phases are separated. (**C**) Two integral membrane proteins (Plp2 and GIRK) and one lipid-anchored protein (Gβγ) tagged with GFP in phase-separated GUVs. The non-raft phase is labeled with a lipid label (either rhodamine-PE or AF594-PE). All three proteins partition exclusively (to the limit of detection) into the non-raft phase, regardless of the percentage of membrane surface area occupied by the non-raft phase (which is controlled by the lipid composition).

| **Fraction of Membrane Protein Signal in Protein-Poor Vesicles** | | | | **Fraction of Membrane Protein Signal in Protein-Rich Vesicles** | | | |
| --- | --- | --- | --- | --- | --- | --- | --- |
| 0.396 | 0.385 | 0.429 | 0.408 | 0.379 | 0.380 | 0.384 | 0.382 |
| 0.404 ± 0.019 | | | | 0.381 ± 0.002 | | | |

**Table S1.** Fraction of membrane protein signal versus total protein signal in protein mass spectrometry data from four independent samples for protein-poor vesicles (left) and protein-rich vesicles (right). The bottom row shows the average and standard deviation for the four samples.
